# APOE4–Aβ synergy drives brain network dysfunction and neuronal lysosomal-ER proteostasis dysregulation a preclinical Alzheimer’s disease model

**DOI:** 10.1101/2025.11.11.687887

**Authors:** Jia Shin, Erica Brady, Chun Chen, Kelli Lauderdale, Ayushi Agrawal, Yutong Zhang, Xueqiao Jiang, Pranav Nambiar, Jessica Herbert, Dakota D. Mallen, Katie K. Ly, Patrick S. Honma, Zhongyi Guo, Cathrine Sant, Reuben Thomas, Stephanie R. Miller, Inma Cobos, Jorge J. Palop

**Author notes:** Contributed equally (Jia Shin, Erica Brady).

## Abstract

Amyloid-β (Aβ) and APOE4 represent two of the strongest pathological and genetic risk factors for Alzheimer’s disease (AD), but how these co-pathogens interact during preclinical stages remains undefined. We addressed this question by developing a humanized knock-in model expressing physiological, endogenously regulated human Aβ and APOE4. Aged *App*^NLF^:APOE4 mice displayed incipient amyloidosis with subtle memory-related changes, consistent with preclinical AD. We found largely distinct, non-overlapping APOE4- and Aβ-driven functional synaptic, sleep, and behavioral alterations. However, at the transcriptomic level, APOE4xAβ had a pronounced detrimental interaction in neuronal populations, whereas glial populations were primarily affected by either genotype. We found APOE4xAβ molecular interactions in neuronal populations, including excitatory and inhibitory cells, converged on a core lysosomal-ER proteostasis axis. We propose that APOE4xAβ interaction produces an early neuronal pathogenic signature, involving the lysosomal-ER proteostasis axis, preceding functional decline and driving disease progression. APOE4xAβ-KI models provide a physiologically relevant platform to study early pathogenesis.

**Highlights:**

- Early synergistic APOE4xAβ interaction emerges predominantly at the transcriptomic level in neurons, but not in glial cells.
- APOE4 and Aβ drive largely non-overlapping physiological changes in preclinical stages of disease, but converge at the level of network hyperexcitability.
- APOE4xAβ neuronal synergy converges on a conserved lysosomal-ER proteostasis axis.
- Humanized APOE4xAβ KI mice provide a physiologically relevant model to dissect early AD pathogenesis in preclinical stages

## Introduction

Alzheimer’s disease (AD) pathology develops over decades before the onset of cognitive symptoms,^1^ with amyloid-β (Aβ) deposition and the APOE4 allele representing the two strongest pathological and genetic risk factors for disease progression. However, as disease progresses, the influence of Aβ deposition^2^ and APOE4^3-5^ becomes notably weaker, particularly after 75 years old, indicating that these two co-pathogens exert their detrimental effects early in disease progression, likely triggering downstream pathogenic cascades during the silent phase of the disease. Yet, the nature of these early interactions and their cell type-specificity remain poorly understood. The preclinical stage of AD is termed asymptomatic amyloidosis (stage 1)^6-14^ and characterized by incipient amyloidosis without tau pathology or overt neurodegeneration^6,12,15-17^. Notably, clinical and imaging studies reveal that both Aβ accumulation and APOE4 produce distinct functional alterations that can precede cognitive symptoms by decades^18-23^. These preclinical alterations are likely driven by distinct Aβ- and APOE4-dependent mechanisms^24-27^, including Aβ-dominant effects on synaptic dysfunction and neural network hyperexcitability, and APOE4-dominant effects on lipids and cholesterol transport, mitochondrial and metabolic stress. However, how APOE4 and Aβ interact under endogenous regulation and physiological expression levels to initiate early molecular changes during incipient amyloidosis and preclinical stages remains unclear.

Recent studies have shown that the convergence of APOE4 and Aβ occurs primarily within glial signaling networks, where APOE4 acts as a critical modulator of both microglial and astrocytic activation states^28-30^. APOE4 promotes the transition of microglia into the disease-associated microglia (DAM) phenotype, characterized by upregulation of TREM2–APOE signaling, enhanced lipid droplet accumulation, and pro-inflammatory transcriptional programs^28^. Aβ deposition and APOE4 expression also drive astrocytic conversion into the disease-associated astrocyte (DAA) phenotype, marked by complement cascade activation, altered cholesterol metabolism, and diminished metabolic and synaptic support to neurons^29,30^. Although these findings highlight APOE4- and Aβ-dependent glial mechanisms, the cell-autonomous neuronal consequences of their interaction before significant amyloid accumulation remain poorly understood.

Contrary to the prolonged silent phase of asymptomatic amyloidosis^6-12^, standard APP transgenic overexpression models with high levels of APP and Aβ expression develop early synaptic dysfunction and dementia-like deficits before the onset of Aβ pathology^31,32^, a disease stage never observed in humans. Thus, these deficits are likely triggered by supraphysiological levels of Aβ/APP expression, which are not found in humans. Instead, humans, even in familial AD (FAD) carriers, show significant amyloidosis preceding cognitive decline, suggesting that increased Aβ production or elevated Aβ_1-42/1–10_ ratios, even for decades, are not enough to cause cognitive impairment in the absence of significant amyloid plaque accumulation. To address these gaps, we developed a humanized knock-in model expressing physiological and endogenously regulated levels of wild-type human Aβ (*App*^NLF^) and human APOE4, avoiding both the pro-amyloidogenic Arctic mutation within the human Aβ sequence present in both *App*^NLGF^ and App^SAA33,34^ and supraphysiological levels of Aβ/APP expression in transgenic models. This model mimics the early stages of AD and functionally balances Aβ- and APOE4-dependent effects with knock-in expression. In aged *App*^NLF^:APOE4 mice, we observed incipient amyloidosis, APOE4- and Aβ-dependent gliosis, mirroring the early glial activation patterns seen in humans with preclinical AD. Despite the absence of overt cognitive impairment, these mice exhibited largely non-overlapping APOE4- and Aβ-dependent deficits in synaptic plasticity, sleep regulation, and spontaneous exploratory behavior. However, at the transcriptomic level, we found pronounced synergistic APOE4xAβ interactions in neurons, including inhibitory and excitatory cells, but not glial cells, whereas glial cells were modulated by either genotype alone with minimal interaction. We found APOE4xAβ neuronal synergy molecularly converged on a lysosomal-ER proteostasis axis. Our study defined early pathogenic APOE4xAβ interactions in neurons during this preclinical stage that may represent early mechanisms driving disease progression.

## Results

### APOE4-driven microgliosis and Aβ-associated astrocytosis in aged humanized E4NLF mice with incipient amyloid pathology

To identify synergistic pathological interactions of APOE4 and Aβ under endogenous regulation and physiological levels during preclinical stages of amyloidosis, we bred human APOE4-KI and App^NLF^-KI mice to obtain APOE4/E4 (E4), *App*^NLF/NLF^ (NLF), APOE4/E4:*App*^NLF/NLF^ (E4NLF), and wild-type (WT) littermates. We selected *App*^NLF/NLF^ mice, instead of *App*^NLGF/NLGF^ or App^SAA/SAA^ mice^33,34^, to study wildtype Aβ without the pro-amyloidogenic Arctic (E693G; G or A) mutation within the Aβ sequence^33,34^. To assess AD-related pathological changes, we quantified Aβ accumulation, microgliosis, and astrocytosis in the hippocampus of aged WT, E4, NLF, and E4NLF mice (14–15-month-old). NLF and E4NLF mice showed incipient levels of Aβ plaques at 14–15 months with NLF mice exhibiting a non-significant trend toward increased amyloidosis compared with E4NLF mice (**Fig. 1a**), since E4 mice have no murine apoE expression^35,36^. Thioflavin-S-positive core mature Aβ plaques were also detected (**Supplemental Fig. 1A**). However, at 18–22 months of age, NLF and E4NLF mice exhibited similar levels of amyloidosis (**Supplemental Fig. 1B–C**). We also compared 14–15-month-old NLF and E4NLF mice with younger (10–13-month-old) *App*^NLGF/NLGF^ (NLGF) mice expressing the additional Arctic mutation within the Aβ sequence (**Fig. 1c**). As expected^34^, NLGF mice had increased amyloidosis compared with NLF and E4NLF mice (**Fig. 1d**). Regarding microgliosis and astrocytosis, we found E4-driven microgliosis in E4 and E4NLF mice (**Figs. 1b and 1e; Supplemental Fig. 1d and 1e**), and NLF-driven astrocytosis in NLF and E4NLF mice (**Fig. 1f**), suggesting a pathogenic divergence between APOE4 and Aβ in driving distinct AD-related pathological changes in early disease pathogenesis, consistent with human transcriptomic data^29,37^. No tau phosphorylation was detected (not shown). These results indicate that aged E4NLF mice have incipient amyloidosis with no tau accumulation but still significant APOE4-dependent microgliosis and Aβ-dependent astrocytosis.

**Fig. 1:**
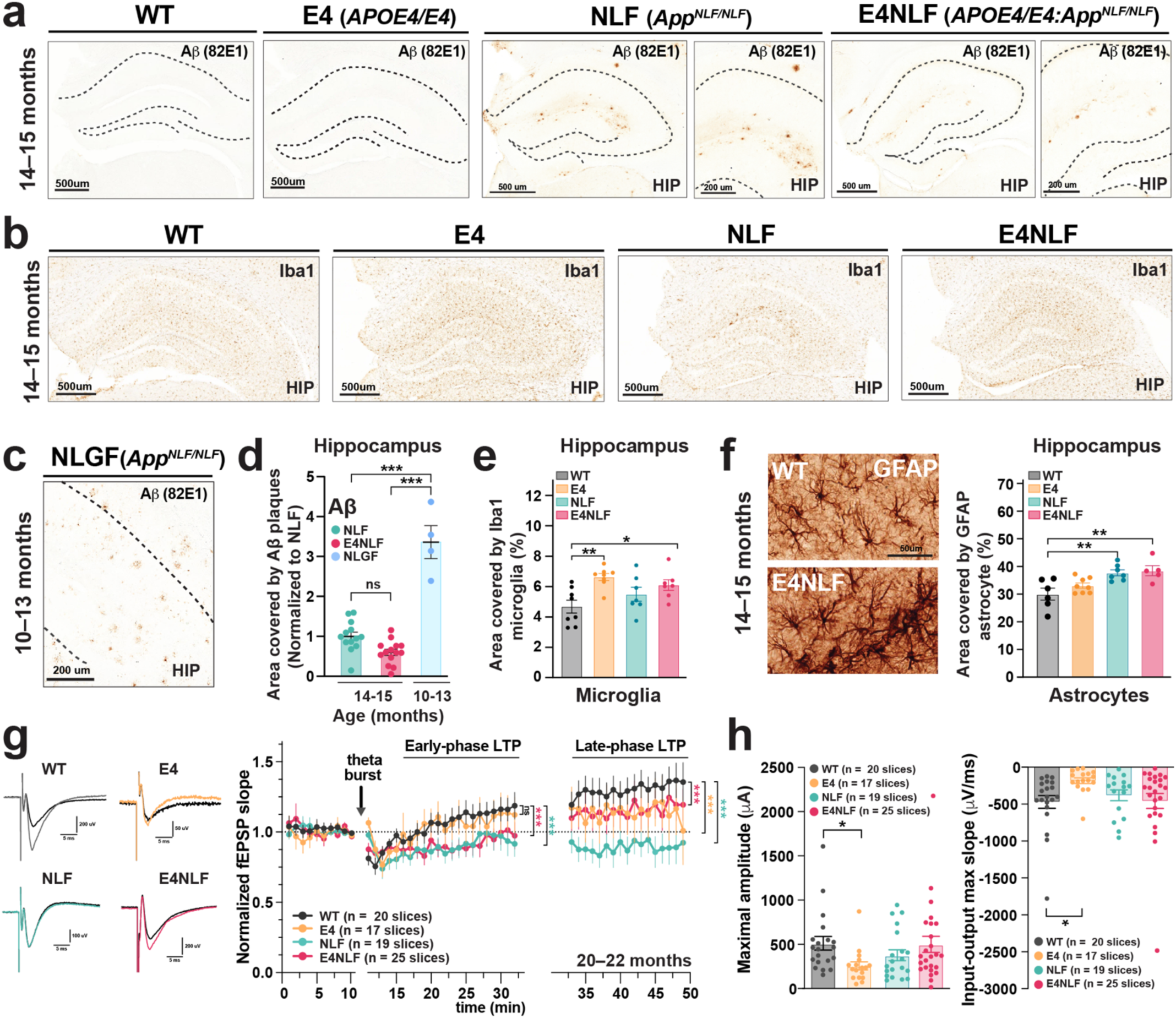
APOE4- and Aβ-dependent synaptic plasticity deficits in aged E4NLF mice with incipient amyloidosis. Aged WT, APOE4/E4 (E4), *App*^NLF/NLF^, and APOE4/E4:*App*^NLF/NLF^ (E4NLF) littermate mice were assessed by electrophysiological and histological analyses. (**a, b**) Representative hippocampal sections immunostained for Aβ (82E1 antibody) (a) and Iba1 (b) in 14–15-month-old WT, E4, NLF, and E4NLF mice. NLF and E4NLF mice displayed incipient amyloidosis and microgliosis consistent with the absence of the pro-amyloidogenic Arctic (E693G) mutation within the Aβ sequence of NLF mice. (**c**) Representative Aβ (82E1) staining in the hippocampus of younger (10–13-month-old) *App*^NLGF/NLGF^ (NLFG) mice with the Arctic (G) mutation showing severe amyloidosis. (**d**) Quantification of hippocampal Aβ accumulation in 14–15-month-old NLF, E4NLF, and NLGF mice. Relative to NLF and E4NLF mutants, NLGF mice had increased amyloidosis due to the pro-amyloidogenic Arctic (E693G) mutation (G). Relative to NLF mutants, E4NLF mice showed a slight reduction in amyloidosis. ***p < 0.001 by one-way ANOVA with Bonferroni post hoc test. (**e**) Quantification of hippocampal microglia in 14–15-month-old WT, E4, NLF, and E4NLF mice. APOE4 induced microgliosis in E4 and E4NLF mice. **p < 0.01, *p < 0.05 by one-way ANOVA with Bonferroni post hoc test. (**f**) Representative GFAP staining in the hippocampus of 14–15-month-old WT and E4NLF mice (left). Quantification of hippocampal GFAP showing increased gliosis in NLF and E4NLF mice (right). **p < 0.01 by one-way ANOVA with Bonferroni post hoc test. (**g**) Field recordings of hippocampal CA1 long-term potentiation (LTP) induced by theta-burst stimulation in 20–22-month-old WT, E4, NLF, and E4NLF mice. Representative field EPSP traces (left) and normalized fEPSP slopes (right) during the early (0–30 min) and late (30–60 min) phase of LTP. Early-phase LTP was impaired by NLF expression, and late-phase LTP was impaired by both E4 and NLF expression compared with WT controls. *p < 0.05, **p < 0.01, and ***p < 0.001 by repeated one-way ANOVA with Bonferroni post hoc test. (**h**) Maximal fEPSP amplitude (left) and input–output maximal slopes (right) revealed deficits in E4 mice. *p < 0.05 by one-way ANOVA with Bonferroni post hoc test. Values are means ± SEM.

### APOE4- and Aβ-dependent synaptic plasticity deficits in aged E4NLF mice with incipient amyloidosis

To determine whether APOE4 and Aβ (wildtype sequence) alter synaptic function under endogenous regulation and physiological levels of expression, we recorded hippocampal CA1 field excitatory postsynaptic potentials (fEPSPs) in aged (20–22-month-old) WT, E4, NLF, and E4NLF mice following theta-burst stimulation. Relative to WT controls, early-phase (0–30 min) long-term potentiation (LTP) was significantly reduced in Aβ-expressing genotypes, including NLF and E4NLF mice, suggesting an Aβ-driven impairment in LTP induction deficits. However, all disease groups, including E4, NLF, and E4NLF mice, showed impaired late-phase (0–30 min) (LTP), suggesting an APOE4-dependent deficit in the maintenance of LTP (**Fig. 1g**). Thus, while Aβ expression attenuated early- and late-phase LTP, APOE4 primarily impaired the late-phase LTP, indicating a potential additive effect on synaptic plasticity. To assess synaptic strength, we analyzed input-output curves revealing decreased maximal fEPSP amplitude and slope in E4 mice, indicating reduced basal synaptic strength (**Fig. 1h**). Together, these results demonstrate that physiological and endogenously regulated levels of APOE4 and Aβ compromise synaptic function and plasticity in aged hippocampal circuits with incipient amyloidosis.

### Aged E4NLF mice develop behavioral alterations but not significant cognitive decline

To determine whether the incipient amyloidosis and gliosis detected in aged E4NLF mice (**Fig. 1**) result in behavioral or cognitive alterations, we tested aged (18–22 months) WT, E4, NLF, and E4NLF littermate mice in the Morris water maze (MWM) and open field to assess cognitive and motor functions. During MWM training, all groups showed progressive improvement in latency to reach the hidden platform, with E4 mice showing subtle differences in the learning curves relative to WT controls (**Fig. 2a**, left). However, 24-h and 72-h probe trials revealed comparable spatial memory retention among genotypes (**Fig. 2a**, right). These results indicate that cognitive performance is largely preserved in this model, which may represent asymptomatic amyloidosis in preclinical AD^6-12^.

**Fig. 2:**
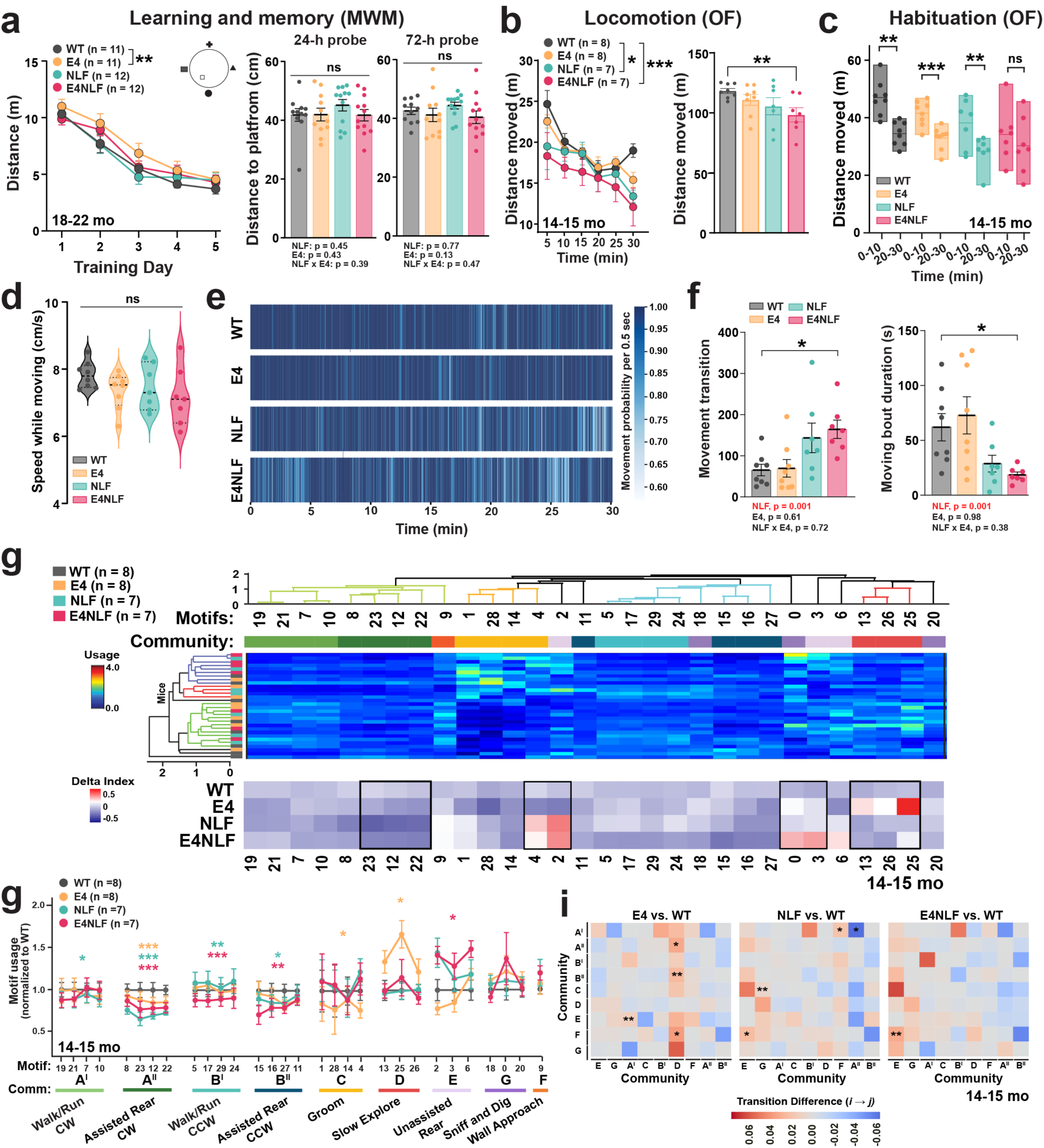
Aged E4NLF mice develop behavioral alterations in spontaneous behavior but not significant cognitive decline. Aged (14–15- and 18–22-month-old) WT, E4, NLF, and E4NLF littermate mice were tested in the Morris water maze, open field, and machine-learning VAME to assess cognitive, motor, and spontaneous behavioral alterations. (**a**) Morris water maze (MWM) learning curves (left) and 24- and 72-hour probe tests (right) for 18–22-month-old WT, E4, NLF, and E4NLF mice. *p < 0.05 by repeated one-way ANOVA with Bonferroni post hoc test. ns by one-way ANOVA with Bonferroni post hoc test. (**b**) Open field (OF) traveled distance in 5-min bins (left) or averaged (right) during 30 min of exploration in an open field in 14–15-month-old WT, E4, NLF, and E4NLF mice. E4NLF mice exhibited reduced locomotor activity compared with WT controls. *p < 0.05, ***p < 0.001 by repeated-measures one-way ANOVA with Bonferroni post hoc test. **p < 0.01 by one-way ANOVA with Bonferroni post hoc test. (**c**) Total distance for the first and last 10 min of open field. All genotypes, except E4NLF mice, showed habituation to the open field. *p < 0.01, **p < 0.001 by paired Student’s t-test. (**d**) Speed during locomotor activity. All genotypes showed comparable speed levels when moving. ns by one-way ANOVA with Bonferroni post hoc test. (**e**) Moving probability (500 ms bins) over the 30 min of open field across all mice and genotypes showing fragmented locomotion sequences and increased immobility in E4NLF mice. (**f**) Moving transition frequencies (left) and moving-bout duration (right) showing increased transitions (more frequent stops) and reduced moving-bout durations in E4NLF mice. **p < 0.01 by one-way ANOVA with Bonferroni post hoc test. Indicated p values for genotypes effects using two-way ANOVA. (**g**) Hierarchical clustering of VAME behavioral motifs (columns) and mice (rows) by motif usage (top). Delta index of motif usage by genotype relative to all genotypes (bottom). (**h**) Relative motif use organized by community across genotypes. E4NLF mice show decreased use exploratory activity, including locomotion and rearing (A^II^, B^I^, B^II^). *p < 0.05, **p < 0.01, ***p < 0.01 by repeated one-way ANOVA with Bonferroni post hoc test. (**i**) Community-level transition matrices comparing E4, NLF, and E4NLF mice with WT controls. Red indicates increased, and blue decreased transition probability relative to WT. E4NLF mice show decreased probability of transitions from exploratory behaviors, including locomotion and rearing (A^II^, B^I^, B^II^) to other behavioral states. *p < 0.05, **p < 0.01 by permutation test. Values are mean ± SEM.

Despite preserved cognitive performance in asymptomatic amyloidosis in humans, behavioral alterations (e.g., apathy, mood symptoms, agitation)^13^ and subtle motor abnormalities (e.g., reduced stride length, slower movement speed, altered gait, rigidity) emerge during this early stage of disease progression, occurring years before the onset of cognitive decline^13,14^. To determine potential motor alterations, we tested 14–15-month-old WT, E4, NLF, and E4NLF mice in the open field. Notably, motor performance was markedly altered in E4NLF mice, displaying a significant reduction in total distance traveled across the 30-min session compared with WT controls (**Fig. 2b**). Interestingly, E4NLF, but not E4 or NLF, mice also showed attenuated habituation between the first and last 10 min of exploration in the open field (**Fig. 2c**), suggesting a subtle learning impairment. To understand the nature of these motor impairments, we assessed locomotor speeds and movement transitions. We found that movement speed during active periods was unchanged among genotypes (**Fig. 2d**), but E4NLF mice instead displayed fragmented locomotor sequences and increased immobility, as indicated by their elevated transition frequencies and reduced moving-bout durations (**Figs. 2e and 2f**). These results indicate that E4NLF mice engage in frequent but shorter episodes of movement, consistent with reduced exploratory drive and apathy-like phenotypes described in early AD^38,39^. Overall, these results indicate a synergistic interaction between APOE4 and Aβ in aged E4NLF mice to trigger subtle motor or motivation-like deficits affecting exploratory behavior.

### Largely non-overlapping E4- and NLF-driven ML-VAME behavioral signatures result in a pathological interaction in E4NLF mice

To determine early signs of dysfunction induced by E4 and Aβ, we applied the ML-VAME platform to segment naturally occurring spontaneous behavior into brief behavioral “motifs” as mice freely explored an open arena^40,41^. Unsupervised motif clustering revealed 30 behavioral motifs organized into 9 communities (A-F), including walk/run, rear, groom, or exploration (**Figs. 2g–I; Supplemental Fig. 2**). Notably, we found a defined behavioral signature in E4, NLF, and E4NLF mice (**Fig. 2g**). Compared to WT, E4 expression induced selective deficits in decreased grooming (C) and increased slow exploration (D), NLF expression induced selective deficits in decreased walk/run (A^I^), and E4 and NLF expression synergized to induce selective impairments in increased unassisted rear (E) and decreased walk/run (B^I^) (**Fig. 2h**). Interestingly, we found only overlapping behavioral deficits between E4 and NLF in decreased assisted rear (A^II^). These results indicate a largely non-overlapping signature of behavioral alterations induced by E4 or NLF expression, but clear signs of synergistic pathological interaction in E4NLF mice. Transition probability matrices further demonstrated reduced transitions from exploratory states to other behavioral communities in E4NLF mice (**Fig. 2i**), highlighting diminished behavioral flexibility and exploratory drive. Collectively, these results demonstrate that aged E4NLF mice exhibit significant alterations in spontaneous and exploratory behaviors despite intact learning and memory performance (**Fig. 2a**), suggesting that the synergistic interaction between APOE4 and Aβ produces behavioral alterations preceding measurable cognitive decline.

### APOE4 and Aβ produce different effects upon sleep architecture but synergistically increase network hyperexcitability

Sleep disturbances are detected during preclinical stages in people at risk of developing AD and are exacerbated by APOE4 genotype^42-44^. To assess APOE4- and Aβ-dependent preclinical changes in the sleep cycle and network hyperexcitability, we performed two-week wireless EEG/EMG recordings in 14–15-month-old WT, E4, NLF, and E4NLF mice in their home cage. Sleep stages were defined using theta and delta power, movement, and EMG activity (see methods), including active wakefulness, quiet wakefulness (QW), non-rapid eye movement (NREM) and rapid eye movement (REM) sleep (**Fig. 3a**)^45^. Notably, compared with WT controls, E4 mice displayed a marked reduction in REM time and REM bouts (**Fig. 3b**), and fewer NREM-to-REM transitions (**Fig. 3b**). REM alterations among genotypes were driven by the APOE4 genotype with milder NLF expression effects. The reduction in REM sleep was most prominent during the light (resting) phase, consistent with human alterations.

**Fig. 3:**
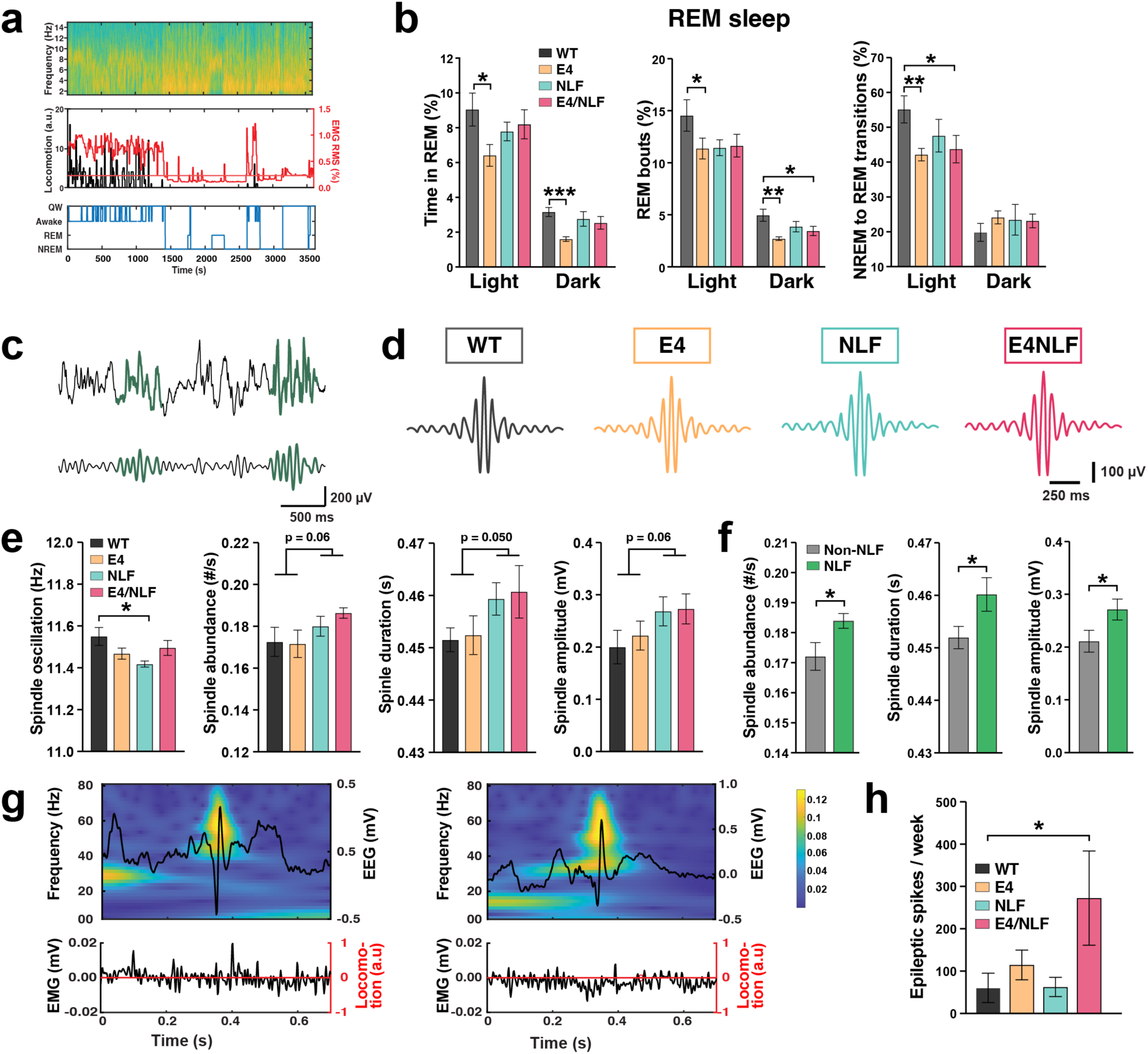
APOE4 and NLF disrupt sleep architecture and synergistically increase subclinical epileptiform activity. Aged (18–22-month-old) WT, E4, NLF, and E4NLF littermate mice were implanted in the posterior parietal cortex with wireless EEG/EMG transmitters and recorded for two weeks to assess sleep architecture and network hyperexcitability. (**a**) Representative EEG spectrogram showing alternating increases in delta and theta power (top), EMG (red) and locomotion (black) traces (middle), and corresponding brain-state classification method of sleep states (bottom), including wake, quiet wakefulness (QW), NREM, and REM. (**b**) Quantification of REM sleep parameters across the circadian cycle, including percent time (left), percent bouts (middle), and NREM–REM transitions. E4 mice show reduced REM time, fewer REM episodes, and decreased transitions to REM compared with WT controls. *p < 0.05, **p < 0.01, ***p < 0.001 by one-way ANOVA with Bonferroni post hoc test. (**c**) Representative raw EEG traces showing characteristic spindle events (10–14 Hz) during NREM sleep. (**d**) Average spindle waveforms across genotypes showing increased amplitude and duration in NLF and E4NLF mice. (**e**) Quantification of spindle oscillation frequency, abundance, duration, and amplitude across genotypes. NLF expression drives spindle alterations in NLF and E4NLF mice. p values two-way ANOVA with NLF and E4 as factors. *p < 0.05 by one-way ANOVA with Bonferroni post hoc test. Indicated p values for NLF genotypes effect using two-way ANOVA. (**f**) Comparison of NLF versus non-NLF genotypes showing that NLF expression drives elevated spindle abundance, duration, and amplitude. *p < 0.05 by unpaired Student t-test. (**g**) Time–frequency spectrograms of representative epileptiform events (top) aligned to locomotor and EMG activity (bottom). E4NLF mice display spontaneous cortical subclinical epileptiform activity. (**h**) Quantification of epileptic spike events for two weeks showing increased synergistic interaction between E4 and NLF expression to trigger cortical hyperexcitability in E4NLF mice. *p < 0.05 by one-way ANOVA with Bonferroni post hoc test. Values are mean ± SEM.

NREM sleep is associated with characteristic thalamocortical spindle oscillations (10–14 Hz) that have been linked to long-term memory consolidation, synaptic homeostasis, and metabolic clearance in humans and mice (**Figs. 3c**)^46,47^. Altered sleep spindle activity has been identified in AD and related models^45,48-50^. Analysis of spindles during NREM sleep revealed NLF-dependent specific alterations in spindle properties (**Fig. 3d–f**), including increased spindle number, duration, and amplitude as well as reduced spindle oscillation frequency. Up and down states during NREM sleep were not affected (**Supplemental Fig. 3**). These findings indicate that Aβ rather than APOE4 primarily drives the disruption of thalamocortical oscillations reflected in sleep spindles.

Subclinical epileptiform activity has been found in asymptomatic APOE4 carriers^51,52^ and early stages of AD^53^, particularly during sleep. Notably, we identified spontaneous epileptiform discharges during sleep (**Fig. 3g**). Epileptic spikes were rare in aged E4 or NLF mice but markedly increased in E4NLF mice (**Fig. 3h**), suggesting a synergistic interaction between APOE4 and Aβ to enhance cortical network excitability. Altogether, these results demonstrate that APOE4 and Aβ exert distinct effects on sleep and network physiology. APOE4 primarily alters REM sleep regulation, while Aβ disrupts thalamocortical spindle dynamics. Their combination synergistically increases cortical hyperexcitability.

### Early synergistic APOE4xAβ interaction emerges predominantly at the transcriptomic level in neurons, but not in glial cells

To identify molecular mechanisms underlying APOE4xAβ interactions at the single-cell level, we performed snRNA-seq analysis of hippocampal tissue from aged (14–15-month-old) WT, E4, NLF, and E4NLF mice (n = 133,413 cells; 6–7 mice per genotype). After quality control-based filtering and clustering, we identified major hippocampal cell types, including interneurons, pyramidal cells, dentate gyrus granule cells, astrocytes, microglia, oligodendrocytes, and OPCs. Pseudo-bulk weighted gene co-expression network analysis (WGCNA) across inhibitory and excitatory pyramidal neurons, microglia and astrocytes revealed pronounced cell- and genotype-dependent shifts in module eigengenes (MEs) (**Fig. 4a**). Notably, in neuronal populations E4 and NLF expression induced largely non-overlapping transcriptional modules, reflecting distinct genetic programs of dysregulation. However, E4NLF mice exhibited distinct synergistic APOE4xAβ alterations not observed in either genotype alone (**Fig. 4a**, left). These results indicate that APOE4 and Aβ each drive independent transcriptomic perturbations, but their co-expression gives rise to a unique molecular signature of disease-associated dysregulation. In contrast, in glial cells, including microglia and astrocytes, NLF and APOE4 expression alone largely drove transcriptional module dysregulation (**Fig. 4a**, right), consistent with the well-documented glial response to amyloid accumulation and APOE4^28-30^.

**Fig. 4:**
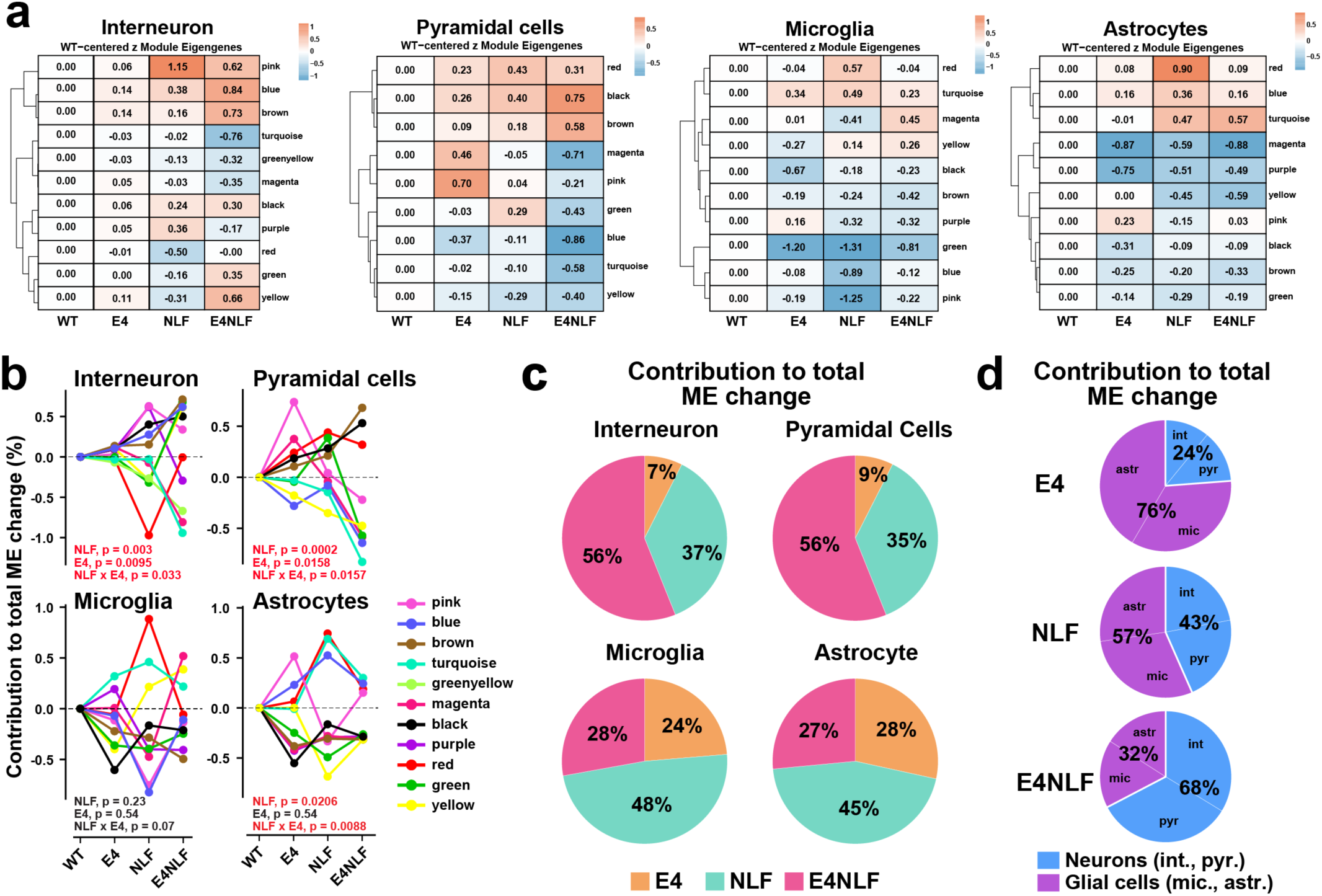
Early synergistic APOE4xAβ interaction emerges predominantly at the transcriptomic level in neurons, but not in glial cells. Hippocampi from aged (14–15-month-old) WT, E4, NLF, and E4NLF littermate mice (n = 7–8 mice per genotype) were dissected and processed for snRNAseq (n = 133,413 nuclei) and canonically clustered into interneurons, pyramidal cells, microglia, and astrocytes. (**a**) Pseudo-bulk weighted gene co-expression network analysis (WGCNA) showing module eigengene (ME) relationships across interneurons, pyramidal cells, microglia, and astrocytes. Module eigengene heatmaps show genotype-dependent relationships for each major hippocampal cell type. (**b**) Percentage contribution to total module eigengene (ME) change across genotype per ME module (total ME change across genotype per module = 1.0). Interneuron and pyramidal cells, but not glial cells, show consistently synergistic interactions between APOE4 and NLF expression in the E4NLF mice. p by two-way ANOVA assessing NLF and E4 genotype and their interaction. (**c**) Percentage contribution to total module eigengene (ME) change by cell type (data from **Fig. 4b**). The contribution of the total ME change in E4NLF mice was higher in interneurons (56%) and pyramidal cells (56%) than in microglia (28%) and astrocytes (27%). (**d**) Percentage contribution to total module eigengene (ME) change by genotype (D) (data from **Fig. 4b**). The contribution of the total ME change in glial cells was higher in E4 (76%) and NLF (57%) than in E4NLF (32%) mice, whereas of the contribution of the total ME change in neurons was higher in E4NLF (68%) than in E4 (24%) and NLF (43%) mice.

To quantify the contribution of ME changes across genotypes and cell types, we calculated the percentage of total module change across genotypes per module (total ME change across genotype per module = 1.0). Neurons, but not glial cells, consistently showed an APOE4xNLF interaction in E4NLF mice (**Fig. 4b**). Pie charts generated from data in Fig. 4B revealed that the contribution of the total ME change in E4NLF mice was driven by changes in interneurons (56%) and pyramidal cells (56%) and to a lesser extend to microglia (28%) and astrocyte (27%) changes (**Fig. 4c**), whereas NLF and E4 changes were predominantly driven by microglia (48% and 24%, respectively) and astrocytes (45% and 28% respectively). Thus, the contribution of the total change in neurons was higher in E4NLF (68%) than in E4 (24%) and NLF (43%) mice, whereas the contribution of total change in glial cells was higher in E4 (76%) and NLF (57%) than in E4NLF (32%) mice. Altogether, our data indicate pronounced synergistic APOE4xAβ interactions in neurons, but not glial cells, whereas glial cells are modulated by either genotype alone with minimal interaction.

### APOE4xAβ interaction alter neuronal gene co-expression networks along a core lysosomal–ER proteostasis axis

To further delineate the biological significance of these APOE4xAβ transcriptional interactions in interneurons and pyramidal cells, we visualized gene expression patterns of the top 50 genes for the top upregulated and downregulated modules for each cell type (**Fig. 5a**) and performed KEGG pathway enrichment analyses (**Fig. 5b**). Notably, both interneurons and pyramidal cells exhibited convergent downregulation of lysosomal and ER proteostasis pathways in their respective modules (magenta and turquoise). Commonly decreased genes included lysosomal and endosomal components (*Ctsb*, *Lamp1*, *Psenen*, *Psene*, *Atp6v0c*, *Tmem30a*, *Tspan3*), ER folding and redox regulators (*Txndc15*, *Itm2c*, *Hspa13*, *Dnajb9*, *C2cd2l*), and vesicle-membrane remodeling factors (*Serinc1*, *Gpm6a*) (**Figs. 5a and 5c**). This coordinated suppression of degradative and folding machineries (ER-lysosomal proteostatic axis) suggests that APOE4xNLF synergy disrupts the neuronal proteostasis network, weakening both the ER quality-control and lysosomal clearance systems required for maintaining protein homeostasis. Similarly, both interneurons and pyramidal cells exhibited shared upregulation of metabolic and cytoskeletal remodeling programs across the blue, yellow, black, and brown modules. Commonly increased genes included those involved in mitochondrial metabolism and oxidative stress resilience (*Acox1*, *Acsf3*, *Nrf1*, *Txnrd2*, *Hspbp1*, *Mettl17*) as well as cytoskeletal and vesicle trafficking regulators (*Bin1*, *Dnm2*, *Camkv*, *Rabep2*, *Map7d1*, *Arhgef25*). These transcriptional adaptations suggest that APOE4xNLF synergy activates conserved neuronal stress responses that enhance energy metabolism and synaptic structural remodeling, likely representing compensatory mechanisms aimed at maintaining cellular stability under pathological pressure. Visualization of shared transcripts confirmed the convergent downregulation of lysosomal genes in E4NLF mice (**Fig. 5d; Supplemental Fig. 4c**). These results reveal that APOE4 and Aβ co-expression coordinately suppress neuronal lysosomal pathways across cortical and hippocampal networks.

**Fig. 5:**
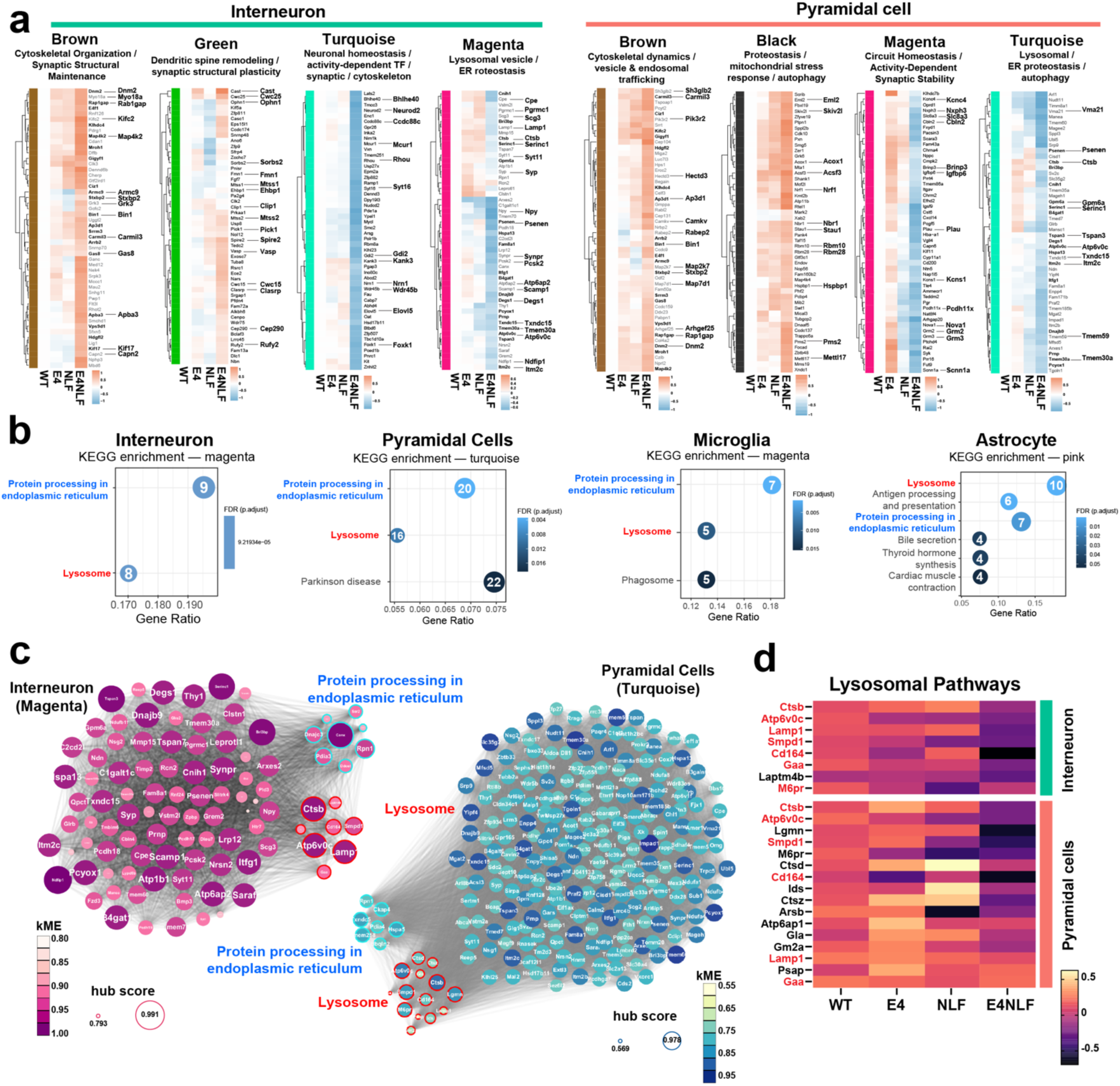
APOE4 and NLF synergistically alter neuronal gene co-expression networks along a core lysosomal–ER proteostasis axis. Hippocampi from aged (14–15-month-old) WT, E4, NLF, and E4NLF littermate mice (n = 7–8 mice per genotype) were dissected and processed for snRNAseq (n = 133,413 nuclei) and canonically clustered into interneurons, pyramidal cells, microglia, and astrocytes. (**a**) Top upregulated and downregulated module-level expression changes in interneuron and pyramidal cells. Top 50 hub genes from each significant module are displayed as heatmaps. (**b**) KEGG pathway enrichment of cell type-specific modules altered in E4NLF mice. Interneurons and pyramidal cells share enrichment in protein processing in the endoplasmic reticulum and lysosomal pathways. (**c**) Network visualization of KEGG-enriched modules for interneurons (magenta) and pyramidal cells (turquoise). Node size represents module connectivity, and color highlights functional annotation. Genes related to lysosomal function (red) and endoplasmic reticulum processing (blue) cluster distinctly, showing coordinated co-expression within each module. (**d**) Heatmap of representative lysosomal genes across genotypes in interneurons and pyramidal cells. E4NLF mice exhibit marked downregulation of lysosomal genes in both pyramidal cells and interneurons, revealing cell type-specific lysosomal dysregulation.

## Discussion

Aβ deposition and APOE4 genotype are, respectively, the most important pathological and genetic risk factors for progressing to AD during its preclinical stages, but the nature of these silent co-pathogenic interactions remains poorly understood. Here, we show that early synergistic pathogenic effects between APOE4 and Aβ during the preclinical stages of AD—with incipient AD-related pathological changes and preserved cognition—emerge predominantly within neuronal populations, including inhibitory interneurons and excitatory neurons, rather than in glial cells, which tend to respond to either pathogen additively. We also found that these pathogenic interactions are primarily detected at the molecular level rather than the functional level. To establish these findings, we developed a novel humanized knock-in model expressing physiological and endogenously regulated levels of wild-type human Aβ (App^NLF^)^34^ and APOE4 (E4NLF)^54^, bypassing ectopic transgenic overexpression and the pro-amyloidogenic mutant Arctic Aβ (E693G), and achieving natural balancing of the expression of these two co-pathogens under endogenous KI regulation. The E4NLF model recapitulates key aspects of the preclinical phase of AD, characterized by incipient amyloidosis at middle-age (15 months) without tau pathology or overt cognitive decline, but subtle memory changes. Notably, APOE4 and Aβ produced largely non-overlapping functional effects with APOE4 preferentially affecting REM sleep and subtle memory-related changes, and Aβ disrupting synaptic plasticity and thalamocortical spindle oscillations. However, robust co-pathogenic synergy between APOE4 and Aβ manifested as enhanced subclinical hyperexcitability during sleep, indicating that hyperactivity may represent one of the first functional drivers of APOE4xAβ interaction in preclinical stages.

At the cellular level, snRNA-seq uncovered a striking neuron-specific synergy between APOE4 and Aβ, while glial cells were largely modulated by either genotype. In interneurons and pyramidal cells, co-expression of APOE4 and Aβ produced unique transcriptional signatures not present in either genotype, marked by the coordinated downregulation of lysosomal and endoplasmic-reticulum (ER) proteostasis pathways and the upregulation of stress-adaptive metabolic and cytoskeletal programs. Together, these results identify a synergistic and convergent neuronal pathway through which APOE4 and Aβ interact under physiological expression levels, potentially triggering downstream pathogenic cascades during the silent phase of AD.

### Modeling the preclinical window of AD in humanized KI mice

AD develops over decades before any cognitive symptoms with progressive Aβ accumulation as the strongest pathological risk factor^6-12^, and APOE4-driven hippocampal hyperactivity^18-21^ as the strongest functional risk factor predisposing individuals to subsequent Aβ accumulation^55^ and memory decline^19^. This preclinical stage of AD, termed asymptomatic amyloidosis (stage 1)^6-14^, holds tremendous promise for early therapeutic interventions. Modeling this prolonged silent phase of asymptomatic amyloidosis has been challenging as standard Aβ/APP transgenic overexpression models consistently develop dementia-like deficits before the onset of Aβ pathology^31,32^—likely due to supraphysiological Aβ/APP expression—a disease stage never observed in either sporadic or in familial AD, where amyloidosis precedes cognitive decline. However, novel humanized Aβ KI models^33,34^ provide physiologically relevant platforms that better recapitulate the early stages of AD pathogenesis, including asymptomatic amyloidosis staging.

We selected the *App*^NLF^ mice, instead of *App*^NLGF^ and *App*^SAA^ mice, to avoid the aggressive pro-amyloidogenic Arctic Aβ mutation (E693G)^56^, which results in amyloidosis by 2–3 months^34^ (<20 years equivalent in humans)^57^, therefore allowing us to study the wildtype human Aβ sequence and incipient Aβ accumulation during aging at 14–15 months of age (>50 years human equivalent)^57^. We crossed the NLF model with the recently developed APOE4-KI mice from the JAX MODEL-AD^54^. Although Aβ/APP transgenic^36,58^ and App^NLGF^ (Ref.)^59^ models have been crossed with APOE4-KI mice, to our best of our knowledge, the E4xNLF genetic interaction has not been previously studied. We found that the E4NLF model isolates physiological APOE4xAβ interactions that mirror asymptomatic amyloidosis staging, including incipient plaques by 14–15 months of age (>50 years equivalent for human)^57^, subtle preclinical memory deficits by 20–22 month (>65 years equivalent for human)^57^, APOE4- and Aβ-dependent gliosis, and preclinical functional abnormalities in sleep and brain hyperactivity. This staging fidelity, along with the endogenously regulated and balanced KI expression of these co-pathogens, is a major strength, particularly assessing transcriptomic changes. A limitation of the E4NLF model is the absence of tau pathology and neurodegeneration. However, this is also consistent with the modeling of the asymptomatic amyloidosis stage where tau is restricted to its normal aggregation patterns in entorhinal and para-hippocampal regions (Braak ≤ 3).

### APOE4- and Aβ-dependent hippocampal hyperactivity and sleep disruptions in preclinical AD

Hippocampal hyperactivity in cognitively normal APOE4 carriers during memory tasks is one of the earliest signatures of AD-related network dysfunction^18-21^. Young (<35 years) and older APOE4 carriers consistently show greater BOLD responses in the hippocampus and medial temporal cortex compared with APOE3 homozygotes during memory tasks, despite equivalent behavioral performance^18-21^. This hyperactivity predicts future cortical Aβ accumulation^55^ and memory decline^19^, suggesting that preclinical APOE4-associated hyperexcitability may be upstream of AD-related pathogenesis. Notably, when assessing Aβ-PiB+ individuals, hippocampal hyperactivity is only observed in MCI (CDR 0.5) subjects, but not in cognitively normal individuals (CDR 0)^22^, suggesting a functional dissociation between APOE4- and Aβ-induced hippocampal hyperactivity. Thus, while APOE4-induced hyperactivation does not require amyloid deposition and cognitive abnormalities, Aβ-induced hippocampal hyperactivity seems to require subtle memory inefficiencies^23^. It is important to note that Aβ-PiB+ cohorts frequently include APOE4 carriers, and most studies do not stratify activation by genotype. To date, there are no published studies that specifically examine hippocampal activation in cognitively normal Aβ-PiB+/APOE3 carriers relative to PiB-/APOE3 carriers. However, there is clear evidence of APOE4-driven hippocampal hyperactivity prior to amyloid deposition at a young age^18,20^. Notably, our longitudinal EEG/EMG analyses revealed that the co-expression of APOE4 and Aβ in E4NLF mice produced emergent hyperactivity, detected as subclinical epileptiform discharges, during NREM sleep, a phenotype absent in NLF but emergent in E4 mice, consistent with the idea that APOE4 establishes a latent hyperexcitable state that is unmasked and exacerbated by Aβ, aligning with a two-hit neural sensitization framework.

Our data strongly parallel the subclinical epileptiform activity in AD patients observed in preclinical APOE4 carriers^51,52^ and early stages of AD^53^, as well as its predominance during NREM sleep and its association with sleep disruptions^60,61^. It also parallels the observed reductions in REM sleep in cognitively unimpaired older adults APOE4 carriers^62^. Although the relationships between hyperactivity, NREM subclinical epileptic activity, sleep disturbances, and APOE4xAβ in older adults have not been disentangled, there is strong evidence for bidirectional causal relationships. For example, Aβ accumulation seems to drive sleep disruptions in healthy older adults and impairs related NREM-sleep-dependent memory consolidation^48^. On the other hand, sleep disturbances are associated with increased Aβ accumulation in older adults^63,64^ and are exacerbated in APOE4 carriers.^65^ In the absence of mechanistic studies in humans, our data suggest that APOE4 may drive sleep disruptions and hyperactivity in the absence of amyloid, but incipient Aβ accumulation may further enhance these functional alterations during preclinical stages.

### Neuronal lysosomal-ER proteostasis dysregulation and associated synaptic dysfunction as a core mechanism of APOE4xAβ interaction in preclinical AD

Our data indicate that the strongest APOE4xAβ interaction emerges in neurons - interneurons and excitatory cells - while glial cells are predominantly modulated by either genotype alone. While prior reports have emphasized Aβ- and APOE4-driven DAM/DAA programs in microglia and astrocytes^28-30^, our data show that incipient amyloid combined with APOE4 triggers an early neuronal response that precedes later stages with more overt glial responses as amyloid accrues and neurodegeneration appears. Notably, APOE4xAβ co-expression without significant AD-related pathology suppresses lysosomal and ER proteostasis pathways, including protein acidification, folding, vesicle trafficking, and degradation. We found many downregulated components of the v-ATPase proton pump, including *Atp6v0c*, *Atp6ap1*, *Atp6ap2*, supporting impaired acidification of lysosomal and synaptic vesicle. Notably, v-ATPase impairments have been linked to lysosomal acidification dysfunction in Down syndrome and AD^66,67^ as well as in APOE4 mice^68^, supporting the proposed framework of early lysosomal dysfunction in AD.^69^ *Lamp1* and Cathepsin B (*Ctsb*) are core components of the lysosomal degradation machinery and were recently identified as critical hub genes of lysosomal dysfunction in AD^70^. Together, these alterations underscore a convergent lysosomal dysfunction in preclinical stages of AD with minimal AD-related pathology.

The endolysosomal system plays a central role in synaptic plasticity by controlling the trafficking, recycling, and degradation of key synaptic receptors, including AMPA and NMDA receptors^68,71,72^. Notably, APOE4 disrupts ApoE receptor–mediated endosomal trafficking, impairing early endosomal maturation and delaying the recycling of glutamatergic receptors and other synaptic cargo, thereby reducing the availability of synaptic receptors required for efficient excitatory transmission and plasticity^71,73^. Our observation of impaired synaptic plasticity in E4 mice is consistent with APOE4- and endolysosomal-deficits in synaptic strength^74^ and plasticity^72,75^ identified in APOE4 mice. Altogether, our data support the notion that early neuronal endolysosomal-ER proteostasis dysregulation and associated synaptic dysfunction disruption represent an early mechanistic event linking genetic and amyloid risk factors in preclinical AD.

## Acknowledgments

We thank the multi-PIs of the PPG project (P01AG073082), Drs. Mucke, Huang, Corces, and Pollard, for their thoughtful input throughout the execution of this project supported by this grant; Drs. Takaomi Saido and Takashi Saito (RIKEN Brain Science Institute) for providing the App^NL-F^ mice. This study was supported by the US National Institutes of Health (NIH) grants R61AG094667, R01AG092683, RF1AG062234, R01AG062629, and P01AG073082 (JJP), and AG082147 (IC), K01-AG083732 (S.R.M.).

## Contributions

Conceptualization: J.S., E.B., J.J.P; Mouse production: E.B., J.H., Y.Z., P.S.H., and J.S.; Histopathology: J.S. and J.J.P.; Electrophysiology: C.C., and Y.Z.; EEG analyses: E.B., J.S., and J.J.P; Standard behavior: J.S., J.H., E.B., and J.J.P.; Machine learning behavior: J.S., K. Ly, S.R.M., P.N., K. Lauderdale., J.H., D.M., and J.J.P; Transcriptomics: J.S., X.J., A.A., R.T., Z.G., C.S., I.C., and J.J.P; Writing: E.B., J.S., and J.J.P. with input from all authors.

## SUPPLEMENTAL FIGURES

**Supplemental Fig. 1 (Related to Fig. 1).**
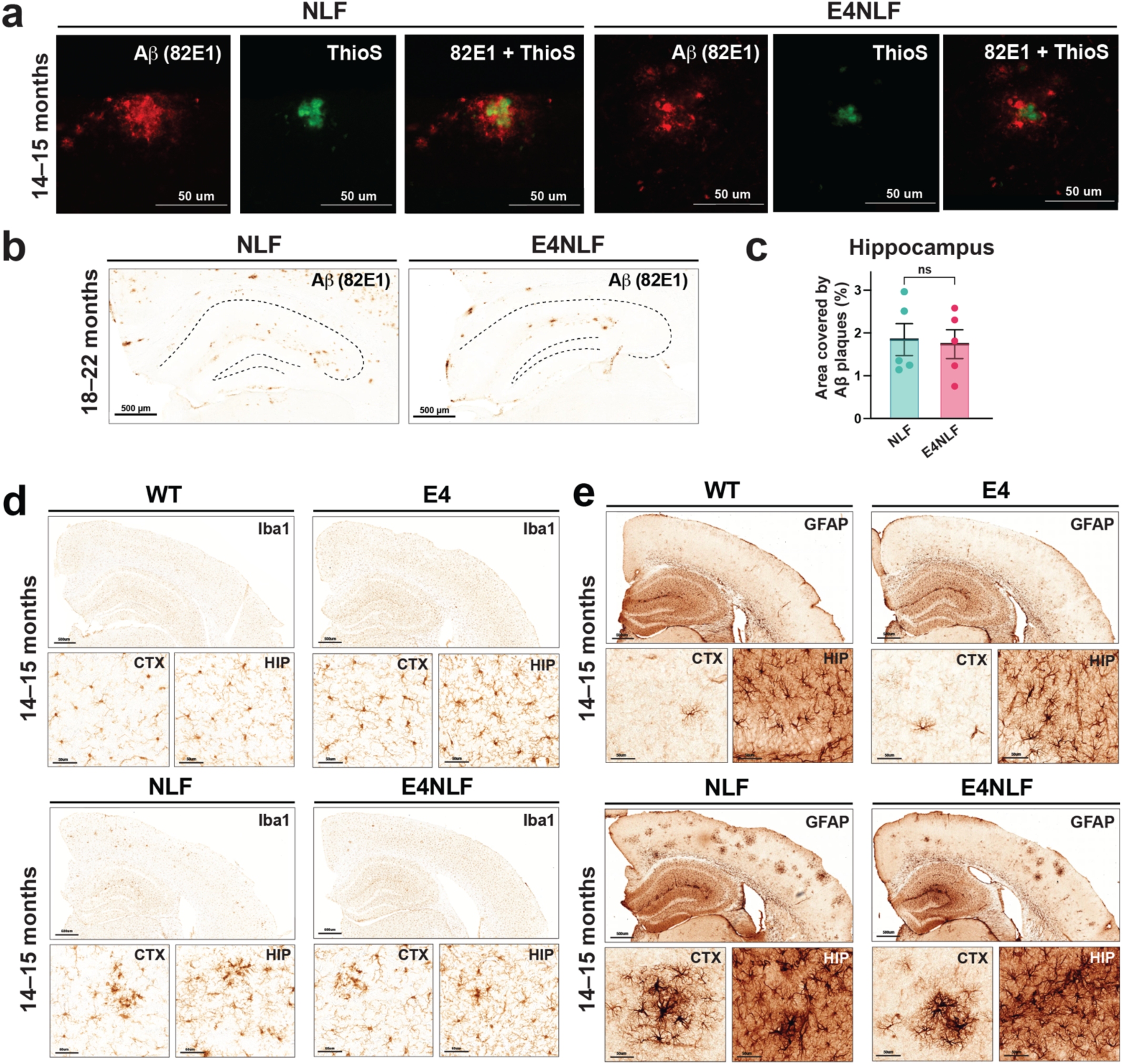
APOE4- and Aβ-dependent pathological changes in aged E4NLF mice. Aged 14–15-month-old (a, d, and e) and 18–22-month-old (b, a) and WT, APOE4/E4 (E4), *App*^NLF/NLF^, and APOE4/E4:*App*^NLF/NLF^ (E4NLF) littermate mice were assessed by histological analyses. (**a**) Representative hippocampal images from 14–15-month-old NLF and E4NLF mice stained with the human-specific Aβ antibody 82E1 (red) and Thioflavin-S (ThioS; green). Mature Aβ plaques in NLF and E4NLF mice display a Thioflavin-S-positive core. (**b**) Representative hippocampal images of Aβ 82E1-immunostained from 18–22-month-old NLF and E4NLF mice. (**c**) Quantification of hippocampal Aβ plaque burden expressed as the percent area covered by 82E1-positive staining in 18–22-month-old NLF and E4NLF mice. No significant differences were observed between 18–22-month-old NLF and E4NLF mice (unpaired two-tailed t-test; ns). Bars represent mean ± SEM. (**d**) Representative hippocampal sections immunostained for Iba1 in 14–15-month-old WT, E4, NLF, and E4NLF mice. Quantification is shown in Fig. 1E. (**e**) Representative hippocampal sections immunostained for GFAP in 14–15-month-old WT, E4, NLF, and E4NLF mice. Quantification is shown in Fig. 1E.

**Supplemental Fig. 2 (Related to Fig. 2).**
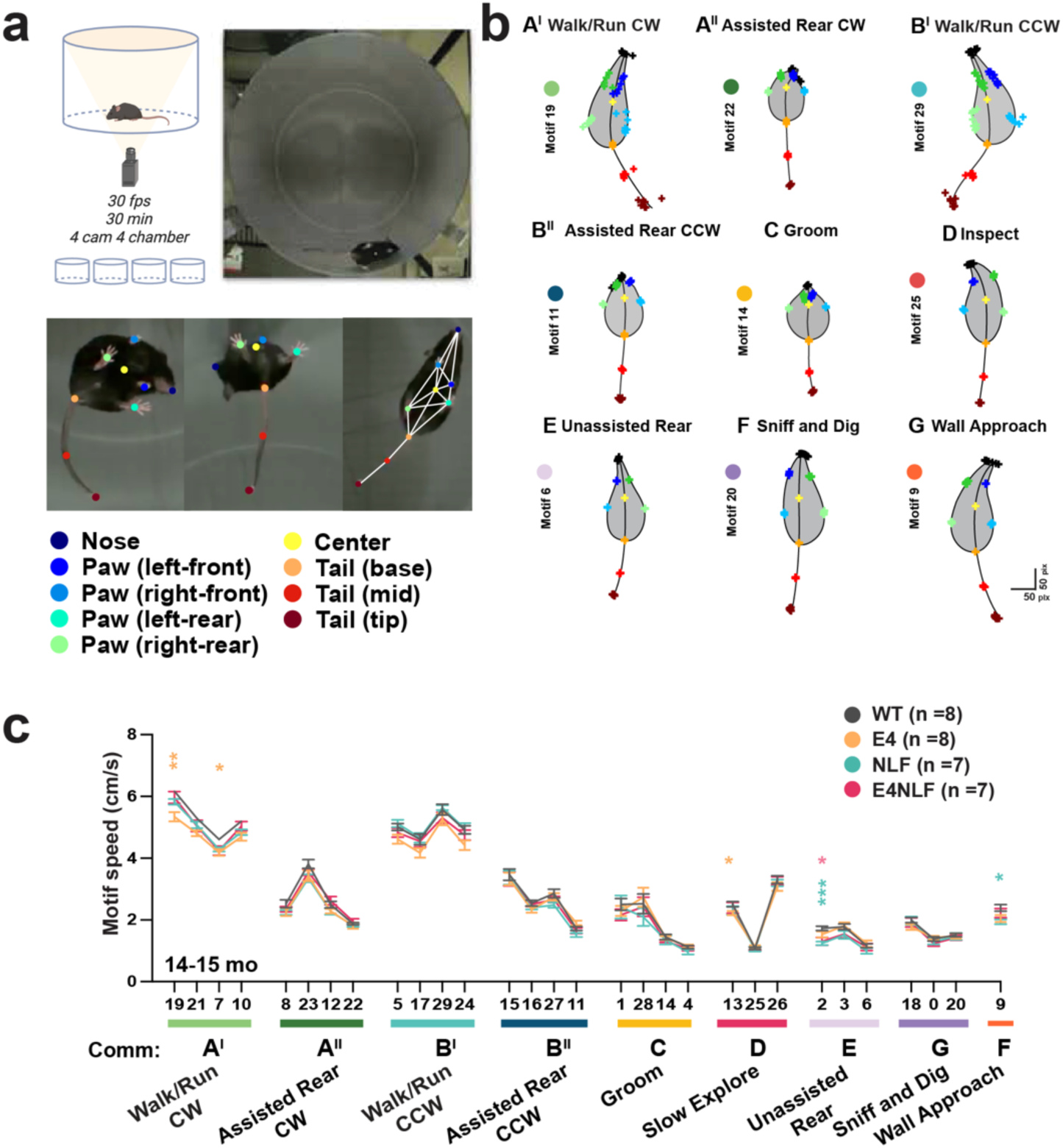
VAME-based unsupervised behavioral segmentation and motif characterization in the open field. Aged (14–15-month-old) WT, E4, NLF, and E4NLF littermate mice were tested using the machine-learning VAME framework to assess spontaneous behavioral alterations. (**a**) Experimental setup and pose-tracking pipeline. Mice were recorded for 30 min in a 4-camera circular open-field arena at 30 fps. Top: schematic of the 4-camera acquisition system and example overhead frame. Bottom: DeepLabCut-based pose estimation using 12 anatomical keypoints (nose, left/right front paws, left/right hind paws, body center, tail base, tail mid, and tail tip) used as inputs for variational animal motion embedding (VAME). Example trajectories and skeletal reconstructions illustrate pose-tracking precision during spontaneous behavior. (**b**) Representative VAME-derived behavioral motifs identified from unsupervised clustering of pose-trajectory embeddings. Each schematic depicts the characteristic posture dynamics and centroid trajectory associated with a specific motif. We identified the following communities: **A′** Walk/Run clockwise (CW): sustained forward locomotion with coordinated paw placement. **A′′** Assisted Rear CW: rear posture supported by the wall with clockwise body rotation. **B′** Walk/Run counterclockwise (CCW): robust locomotion with CCW trajectory curvature. **B′′** Assisted Rear CCW: rear posture supported by the wall with CCW rotation. **C** Groom: cyclical head–body movements characteristic of self-grooming. **D** Inspect: stationary exploratory posture with forward head extension. **E** Unassisted Rear: vertical rearing without wall support. **F** Sniff and Dig: nose-down exploratory sequences with paw and snout movements. **G** Wall Approach: directed movement toward the arena perimeter followed by orienting behavior. Scale bar, 50 px (approx.). (**c**) Motif-specific speed across genotypes. Line plots show average centroid speed (cm/s) for each motif, aligned to the corresponding motifs and communities. Motif speeds were comparable across genotypes, with modest differences limited to a subset of high-speed locomotor motifs (two-way ANOVA with Bonferroni post hoc tests; *p* < 0.05 where indicated). Data represent mean ± SEM.

**Supplemental Fig. 3 (Related to Fig. 3).**
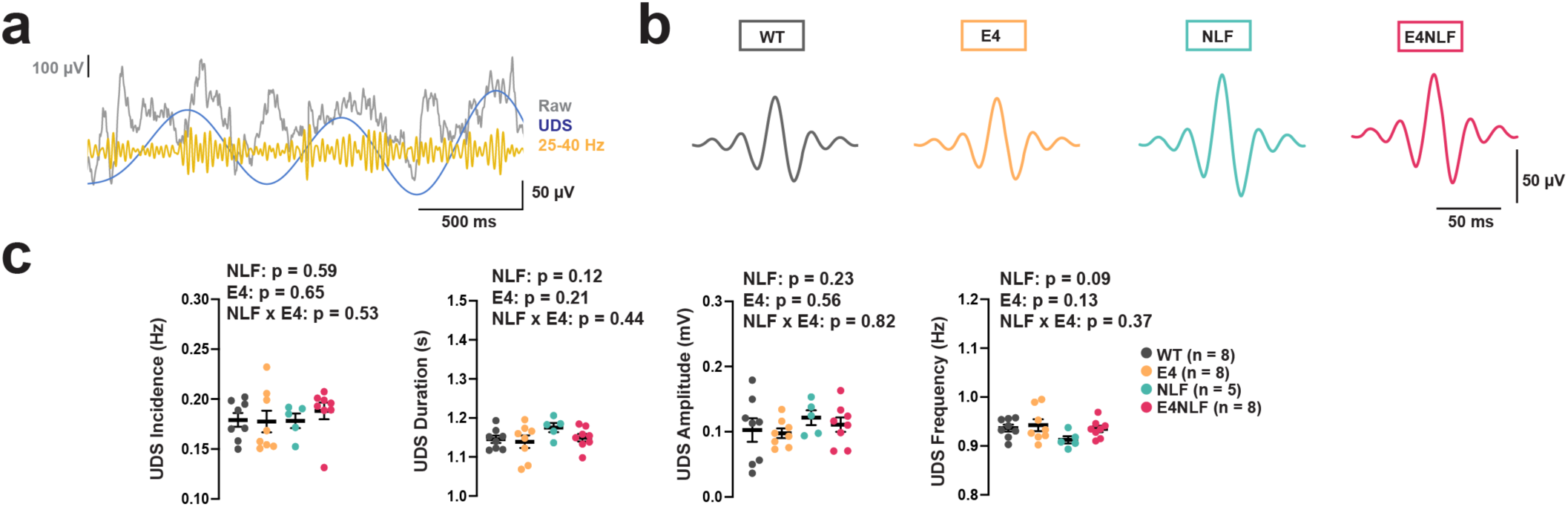
Quantification of cortical Up–Down state (UDS) dynamics across genotypes. Aged (18–22-month-old) WT, E4, NLF, and E4NLF littermate mice (WT n = 8, E4 n = 8, NLF n = 5, E4NLF n = 8) were implanted in the posterior parietal cortex with wireless EEG/EMG transmitters and recorded for two weeks to assess slow-oscillation dynamics during NREM sleep. (**a**) Representative raw EEG trace (grey) with band-pass filtered 25–40 Hz envelope (yellow) illustrating alternating Up and Down states detected during NREM sleep. Blue line shows the algorithm-derived UDS segmentation. (**b**) Average Up–Down state waveforms across genotypes, aligned to Up-state onset. WT (black) displays a canonical depolarizing Up state followed by a hyperpolarizing Down state; E4 (orange), NLF (green), and E4NLF (magenta) mice show comparable waveform morphology. (**c**) Quantification of UDS parameters, including incidence (Hz), duration (s), amplitude (µV), and oscillation frequency (Hz). Across all measures, no significant APOE4-, NLF-, or APOE4×NLF-interaction effects were detected (two-way ANOVA with NLF and E4 as factors; indicated p values). Scatter plots show individual mice and group means ± SEM.

## Methods

### Animals

App^NL-F^ mice were obtained from the RIKEN Brain Science Institute (Drs. Takaomi Saido and Takashi Saito, Japan)^1^. App^NL-F^ mice knock-in express humanized Aβ with the familial Alzheimer’s disease (FAD) Swedish (KM670/671NL) and Iberian (I716F) mutations, but not the aggressive pro-amyloidogenic Arctic Aβ mutation (E693G), on a C57BL/6J background. Human APOE4 knock-in mice, in which the murine Apoe gene is replaced with the human APOE4 allele, were obtained from The Jackson Laboratory (C57BL/6J background; JAX MODEL-AD)^2^. To generate the double knock-in cohort (APOE4; App^NL-F^), we first crossed homozygous APOE4 and homozygous App^NL-F^ mice to produce F1 heterozygotes, which were intercrossed to yield all four homozygous genotypes in the F3 generation: wild-type (WT), APOE4/E4 (E4), App^NLF/NLF^ (NLF), APOE4/E4:*App*^NLF/NLF^ (E4NLF) littermates mice. Both male and female homozygous animals were used for experiments as indicated in the experimental mouse cohort section below. Mice were group-housed under standard conditions with ad libitum access to food and water on a 12-hour light/dark cycle. All mice were bred and maintained in the Gladstone Institutes vivarium (UCSF), an AAALAC-accredited facility. All procedures were approved by the UCSF Institutional Animal Care and Use Committee and conducted in accordance with NIH guidelines for the care and use of laboratory animals.

### Experimental mouse cohorts and sexes

Open field was performed in 14–15-month-old WT, E4, NLF, and E4NLF mice (male), including WT (n = 8, mean age = 14.71 ± 0.71 months), E4 (n = 8, mean = 15.29 ± 0.24), NLF (n = 7, mean = 15.23 ± 0.00), and E4NLF (n = 7, mean = 15.20 ± 0.25).

Morris water maze was performed in 20–22-month-old WT, E4, NLF, and E4NLF mice (female), including WT (n = 11, mean age = 20.08 ± 0.60 months), E4 (n = 11, mean = 20.01 ± 0.70 months), NLF (n = 12, mean = 20.28 ± 0.70 months), and E4NLF (n = 12, mean = 19.83 ± 0.91 months).

ECoG recordings were performed in 18–22-month-old female WT, E4, NLF, and E4NLF mice, including WT (n = 10, mean age = 19.87 ± 0.77 months), E4 (n = 9, mean = 20.80 ± 1.13 months), NLF (n = 9, mean = 20.12 ± 1.27 months), and E4NLF (n = 7, mean = 20.36 ± 1.62 months).

Slice physiology was in 20–22-month-old WT, E4, NLF, and E4NLF mice (female), WT (n = 4, mean age = 22.16 ± 2.19 months), E4 (n = 3, mean = 22.54 ± 4.68 months), NLF (n = 4, mean = 21.79 ± 2.08 months), and E4NLF (n = 4, mean = 21.18 ± 2.46 months).

We used only females for assessing cognition and physiology because it has been reported that they are more susceptible to APOE4-induced deficits^3-5^. Males and females were used for behavior and histopathology. We found no significant histopathological differences between males and females (not shown). Males were used for scRNAseq analyses. Experimenters were unaware of the genotype when conducting the experiments and collecting the data.

### Behavioral experiments

#### Morris water maze

A circular pool (122 cm diameter) contained opaque water, and a square escape platform (14 cm²) was submerged 1 cm below the surface. Fixed distal cues surrounded the pool. Mice completed 10 hidden-platform sessions across five days, two sessions per day spaced 3–4 h apart. Each session had two 60 s trials with a 15 min intertrial interval. Start locations varied pseudorandomly while the platform position was constant. Twenty-four hours after the last session, a 60 s probe trial was run with the platform removed. A visible-platform control followed at the same location, marked by a pole. Behavior was tracked in EthoVision XT, and latency, path length, and swim speed were recorded. Animals that failed the visible-platform task and exceeded their genotype variability (>3 SD) were excluded.

#### Open field

Spontaneous activity was assessed in a circular acrylic arena (1 m diameter, 40 cm high) lined with laminated white paper to minimize visual cues. Mice underwent a 1-hour light habituation period in the testing room prior to the experiment. They were then placed in the center of the arena and allowed to explore freely for 30 min during the light phase. Trials were randomized across genotypes to control for circadian effects. Behavior was recorded at 30 fps using an overhead camera (Spinview) under dim lighting (1% brightness, 5600K). The arena was cleaned with 70% ethanol between sessions.

##### Movement state analysis

EthoVision® XT (Noldus, RRID:SCR_000441; RRID:SCR_004074) was used to track center-point movement and classify locomotor states in 0.5 s bins. Each bin was assigned a binary state (1 = moving, 0 = not moving) based on whether displacement exceeded the movement threshold. Locomotor probability per bin was averaged across animals within genotype to generate time-resolved movement trajectories across the 30-minute session, enabling visualization of genotype-specific exploratory dynamics.

##### Moving bout analysis

To quantify the structure of spontaneous locomotion, consecutive bins with identical states (moving or not moving) were grouped into movement bouts and stillness bouts. For each mouse, we quantified (i) the number of bouts in each state, (ii) the number of transitions between states, and (iii) the mean duration of each bout type (total time in state / number of bouts). Mean movement speed was calculated only within moving bouts (total distance / total movement duration). These metrics capture behavioral persistence versus fragmentation, allowing genotype-level comparison of locomotor organization.

#### VAME-based unsupervised behavioral segmentation and motif characterization

We adopted the VAME-LINC (v1.0) framework developed by Luxem et al.^6^ and Miller et al.^7^ for unsupervised behavioral segmentation of pose data derived from DeepLabCut (v2.3.9). DeepLabCut was used to tracking the finer grain postures by labeling nine body parts (nose, left/right forepaws, left/right hindpaws, belly, tail base, mid-tail, and tail tip) using a ResNet-50 backbone pretrained on ImageNet. A training set was constructed by extracting 10 representative frames per video from a sex- and genotype-balanced subset of 25 open field recordings, with all frames manually annotated. The network was trained for up to 634,000 iterations, reaching a training error of 1.11 pixels (0.82 mm) and a test error of 4.42 pixels (3.28 mm), indicating high accuracy. Pose data were then egocentrically aligned using the VAME alignment module, anchoring the body axis along the nose–tail base vector. Mid-tail and tail tip coordinates were excluded from the VAME input due to their delayed motion. All 30 mice were processed under a single VAME-LINC model trained with default settings: 30 latent dimensions (zdims), a 12-frame time window (time_window), and 30 behavioral clusters (n_clusters) using the standard RNN architecture. After convergence, behavioral motifs were assigned to each frame using a 16-frame sliding window (±320 ms), and final clustering was performed using k-means.

#### Behavioral motif clustering and validation

All final motif assignments were based on k-means clustering. Annotated motif-labeled and community-labeled videos were generated for each cohort to enable visual inspection. VAME-LINC outputs were exported and processed further in a custom Google Colab notebook (see Code Availability).

##### Motif usage correlation cluster analyses

To examine similarity in motif usage across animals/genotypes and identify clusters of commonly used motifs, correlation clustering was applied to the motif usage matrix (rows = mice, columns = motifs). Pearson’s correlation was calculated between each subject’s motif usage profile and the WT group-averaged profile, and subtracted each element of the resultant correlation matrix from 1 to obtain the dissimilarity matrix. MATLAB’s linkage() and dendrogram() functions were used to generate a correlation cluster tree, grouping motifs and subjects with similar usage patterns. Clustering was performed along both dimensions, enabling reordering of the matrix based on the output permutation from dendrogram(). This approach allowed for joint visualization and identification of motif and subject clusters based on similarity in motif usage patterns.

##### Motif characterization and community grouping

Behavioral motifs extracted from the VAME pipeline were annotated through a blinded, multi-reviewer process. Each reviewer analyzed five sessions with balanced genotype representation and partial overlap to ensure labeling consistency. Motifs were described using standardized behavioral terms, referencing fine kinematics and noting any distinctive or anomalous patterns. Following individual reviews, a Motif Consensus Meeting was held to finalize primary and secondary labels based on group discussion. Motifs were then grouped into behavioral communities by clustering similar annotations, guided by correlation-based motif usage dendrograms as well as pose estimation and motif-specific speeds. Final community groupings and semantic labels were refined through collaborative discussion and recorded by the Lead Analyst for downstream analyses.

##### Motif reprocessing for visualization

Initial motif clustering for this cohort was performed using VAME-LINC (v1.0) with 6 body points, excluding the tail base, mid-tail, and tail tip, in order to expedite processing. To generate improved egocentric visualizations for figure representation, the same dataset was reprocessed using VAME v0.5.1 with an expanded set of body points including the tail base, mid-tail, and tail tip. This reprocessing was performed in a separate VAME project and did not alter the original motif assignments, but served to enhance the kinematic depiction of representative motifs for characterization and presentation purposes.

##### Community glossary

We identified the following communities:

**A**′ Walk/Run clockwise (CW): sustained forward locomotion with coordinated paw placement.

**A**′′ Assisted Rear CW: rear posture supported by the wall with clockwise body rotation.

**B**′ Walk/Run counterclockwise (CCW): robust locomotion with CCW trajectory curvature. **B**′′ Assisted Rear CCW: rear posture supported by the wall with CCW rotation.

**C** Groom: cyclical head–body movements characteristic of self-grooming.

**D** Inspect: stationary exploratory posture with forward head extension.

**E** Unassisted Rear: vertical rearing without wall support.

**F** Sniff and Dig: nose-down exploratory sequences with paw and snout movements.

**G** Wall Approach: directed movement toward the arena perimeter followed by orienting behavior.

#### Community transition matrix

Motif transition matrices were derived from VAME motif sequences. First, a motif-level transition matrix was computed for each session by counting transitions between motifs and row-normalizing to control for total activity; self-transitions (motif → same motif) were excluded, as remaining in the same motif is trivially 1. Motifs were then mapped to one of nine behaviorally defined communities, and community-level transition probabilities were obtained by averaging transition values across all motif pairs belonging to each community transition, again excluding self-transitions at the community level. Because the animal is the true unit of replication, session-level matrices were averaged within animal prior to group-level comparisons.

Genotype effects were quantified using Δ-transition matrices, computed as (genotype − WT) for each from→to community transition, such that positive values indicate increased transition likelihood relative to WT. To assess whether Δ exceeded chance variation, we performed a non-parametric animal-block permutation test, in which genotype labels were randomly shuffled at the animal level while preserving the number of animals per group. This approach generates a null distribution for Δ without assuming normality or equal variance. Two-sided permutation p-values were computed by comparing observed Δ to this null distribution. Results are visualized as zero-centered heatmaps, where color indicates the magnitude and direction of Δ, and statistically significant transitions are marked.

#### Histology and immunohistochemistry

##### Tissue preparation

All mice were anesthetized and transcardially perfused with nuclease-free 0.1 M phosphate-buffered saline (NF-PBS). Brains were extracted and drop-fixed in 4% phosphate-buffered paraformaldehyde at 4 °C for 48 hours. After rinsing with PBS, tissue was cryoprotected in 30% sucrose in PBS at 4 °C for 24 hours, coronally sectioned at 40 µm on a sliding microtome, and stored at −20 °C in cryoprotectant solution. Prior to staining, free-floating sections were washed in PBS and permeabilized in 0.5% PBST.

##### DAB immunohistochemistry

Endogenous peroxidase activity was quenched in 3% H₂O₂ in methanol. Sections were blocked in 10% normal donkey serum with 0.2% dry milk and 0.2% gelatin in PBST.DAB Immunohistochemistry. Cryosectioned brain sections were dried, post-fixed (if unfixed), and incubated overnight at 4°C with primary antibodies against Aβ, Iba1, GFAP, c-Fos, or PSD-95. After PBS-T washes, sections were incubated with biotinylated secondary antibodies (for non-biotinylated primaries), treated with the VECTASTAIN ABC-HRP Kit (Vector Laboratories, Cat# PK-6100) and visualization with DAB Peroxidase Substrate (Vector Laboratories, Cat# SK-4100). Slides were dehydrated, cleared in xylene, and cover slipped.

##### Primary antibodies

Aβ (82E1), rabbit anti-Aβ (1:500; IBL #18584, RRID: AB_10705431); Aβ (82E1), mouse anti-Aβ, biotinylated (1:500; IBL #10326, RRID: AB_10705565); Iba1, microglia marker (rabbit anti-Iba1, 1:1000; FUJIFILM Wako #019-19741, RRID: AB_2314669 / AB_228212); GFAP (mouse anti-GFAP, 1:3000; Millipore Sigma, Cat# MAB360).

##### Secondary antibodies

biotinylated anti-rabbit IgG (Invitrogen #31821, RRID: AB_228212); Alexa Fluor 594 goat anti-rabbit IgG (1:1000; Thermo Fisher Scientific, Cat# A-11012, RRID: AB_2534079); biotinylated goat anti-rabbit IgG (1:500; Invitrogen, Cat# 31821, RRID: AB_228212).

##### Fluorescence double staining

For fluorescence detection of Aβ plaques, sections were incubated overnight at 4 °C with rabbit anti-Aβ (82E1) (IBL #18584, RRID: AB_10705431), followed by Alexa Fluor 594-conjugated anti-rabbit IgG (Thermo Fisher, see catalog number used in your lab). Sections were subsequently stained in 0.5% Thioflavin S in ethanol, rinsed, and mounted in antifade medium containing DAPI.

##### Imaging and quantification

DAB-stained sections (Aβ, Iba1, and GFAP) were imaged on a Leica Versa 200 Slide Scanner under uniform exposure settings. Fluorescent Aβ/Thioflavin S–labeled sections were imaged on an Evident FV3000RS confocal microscope with consistent laser intensities and PMT gain. Plaque burden was quantified in ImageJ. Anatomical ROIs were defined using the Allen Mouse Brain Atlas.

#### Surgical electrocorticography (ECoG) implantation and data acquisition

Mice were induced with isoflurane in oxygen (3%) and maintained at 0.5–1% throughout surgery. After shaving, animals received buprenorphine (0.05 mg/kg, i.p.) and were secured in a stereotaxic frame. The scalp was cleaned with alternating 70% ethanol and povidone-iodine. Lidocaine was applied locally, then a midline incision was made and extended posteriorly with a slight angle toward the left shoulder. A subcutaneous pocket was formed along the left dorsolateral flank and filled with about 500 µL sterile 0.9% sodium chloride (bacteriostatic). A sterilized HD-X02 telemetry transmitter (∼2 g; Data Sciences International) was placed subcutaneously on the left side. After clearing periosteum, two burr holes (∼1–2 mm) were drilled at roughly −2 mm AP and ±2 mm ML from bregma. ECoG leads were de-insulated at the tips, gently looped, and slid just beneath the skull over dura to contour the posterior parietal and somatosensory regions. The loops were seated under the skull, the free ends left external, and dental acrylic was used to secure the assembly. For EMG, insulation was removed from the leads and the wires were slightly tensioned to length. A sterile 25-gauge needle guided the leads into a right-side neck muscle, placing the two wires about 3–4 mm apart within the same muscle group. Sutures fixed the wires in place and excess exposed length was trimmed. The incision was closed with sutures. Postoperative analgesia included buprenorphine (0.05 mg/kg) and ketoprofen (5 mg/kg). Animals recovered in warmed, clean cages under continuous observation until ambulatory, then returned to home cages and checked daily for 3–4 days. ECoG /EMG recordings began no earlier than day 7 to allow full recovery and stabilization of the implant.

##### Data acquisition and synchronization

Wireless ECoG and EMG were recorded continuously for 14 days with the DSI Ponemah system at 500 Hz. Infrared locomotor tracking (ANY-maze) ran in parallel and was time-aligned to annotate behavioral state such as rest versus active movement. Recordings took place in sound-attenuated, temperature-controlled chambers on a 12 h light/dark cycle to limit environmental and circadian variability.

#### Slice physiology and long-term potentiation (LTP) methods

LTP experiments were conducted on acutely prepared coronal hippocampal slices. Slice preparation followed the protective recovery method described previously (citation: Jonathan Ting et al.). Mice were anesthetized with avertin (tribromoethanol; 0.15 mL per 10 g body weight) and transcardially perfused with ice-cold, carbogenated (95 % O₂/5 % CO₂) NMDG-based dissection solution containing (in mM): 93 N-methyl-D-glucamine, 2.5 KCl, 1.2 NaH₂PO₄, 30 NaHCO₃, 20 HEPES, 25 D-(+)-glucose, 5 ascorbic acid, 2 thiourea, 3 sodium pyruvate, 12 N-acetyl-L-cysteine, 10 MgSO₄, and 0.5 CaCl₂, adjusted to pH 7.3–7.4. Following decapitation, brains were rapidly removed and coronal sections (300 μm) containing the hippocampus were cut on a vibrating microtome and transferred to an incubation chamber containing warmed (35 °C) dissection solution. Sodium concentration was gradually increased depending on the age of the animals until it matched that of the recovery ACSF containing (in mM): 92 NaCl, 2.5 KCl, 1.2 NaH₂PO₄, 30 NaHCO₃, 20 HEPES, 25 glucose, 5 ascorbic acid, 2 thiourea, 3 sodium pyruvate, 12 N-acetyl-L-cysteine, 2 MgSO₄, and 2 CaCl₂. Slices were then transferred to a holding chamber containing oxygenated recovery ACSF at room temperature for at least 1 h prior to recording.

Extracellular field recordings were performed using the MED64 Quad II system (Alpha MED Scientific) equipped with four 4 × 4 electrode arrays (PG515A; 50 μm electrode diameter, 150 μm inter-electrode spacing). Arrays were pre-coated with polyethylenimine to improve tissue adhesion. Slices were positioned so that the CA1 region overlaid the array, aligning the Schaffer collateral projection approximately along the array electrode region. Slices were continuously perfused with oxygenated ACSF, containing (in mM): 119 NaCl, 2.5 KCl, 1 NaH_2_PO_4_, 26.2 NaHCO_3_, 11 Glucose, 1.3 MgSO_4_, 2.5 CaCl_2_ at 2-3 mL/min and maintained at 32 °C throughout the experiment.

LTP induction, Theta-burst stimulation (TBS), was delivered through a single stimulation electrode placed in the stratum radiatum. A recording electrode located 150 μm (single electrode spacing) downstream along the Schaffer collateral pathway was selected. An input–output (I/O) curve was generated at the beginning of each experiment to determine the stimulation intensity required to elicit the maximal fEPSP response. Biphasic constant-voltage pulses (0.2 ms) were delivered starting from 0 µV, and the stimulus intensity was increased in 5 µV increments at 20 s intervals. The maximal fEPSP amplitude was defined as the plateau of the I/O curve, and 30–50 % of this maximal response was used as the stimulation intensity for baseline and LTP recordings. Baseline fEPSPs were recorded every 20 s for 10 min before induction and 40 min post induction. TBS consisted of 5 trains at 20 second interval, each containing 10 bursts delivered at 10 Hz. Each burst comprised 4 pulses at 100 Hz. The stimulus intensity was set to 70 % of the maximal response to reliably engage synaptic plasticity while minimizing excessive population spike activity. The slope of the fEPSP was measured on the rising phase between 10–40 % of the peak amplitude. LTP was expressed as the percentage change in the mean fEPSP slope and amplitude.

#### scRNAseq analyses

##### Tissue preparation, fixation, and nuclei isolation

Hippocampal tissue was dissected from flash-frozen coronal brain sections stored at −80 °C. Tissue was immediately transferred to ice-cold PBS and mechanically homogenized using loose and tight Dounce pestles in Nuclei Isolation Buffer (10x Genomics, Chromium Fixed RNA Profiling Kit; CG000478) supplemented with 1% RNase inhibitor. The homogenate was filtered through a 40 μm strainer and centrifuged at 500 × g for 5 min at 4 °C. The nuclear pellet was resuspended in Fixation Buffer and incubated at room temperature for 30 min according to the manufacturer’s instructions to preserve RNA integrity and transcript abundance patterns. Following fixation, nuclei were washed twice in Wash Buffer and resuspended in Storage Buffer at 4 °C prior to downstream processing. All steps were performed on ice unless otherwise noted to maintain nuclear integrity and prevent RNA degradation.

##### Library Preparation and sequencing

Fixed nuclei were counted and diluted to the recommended input concentration (700–1,200 nuclei/μL), and ∼6,000 nuclei were loaded per reaction onto a Chromium X controller (10x Genomics). Single-nucleus barcoding and library construction were performed using the Chromium Fixed RNA Profiling Reagent Kit and workflow (10x Genomics, CG000478) following the manufacturer’s protocol. Libraries were sequenced on an Illumina NextSeq platform at the Gladstone Institutes Genomics Core at a target depth of ∼50,000 read pairs per nucleus. A total of 30 libraries were generated, of which one (sample 993) was excluded due to poor quality metrics in Cell Ranger output

##### Processing of Single-Nucleus RNA-Seq Data

Demultiplexed FASTQ files were processed using Cell Ranger v7.2.0^8^ with the 10x Genomics pre-built mm10 reference and the include-introns option enabled. Gene-barcode matrices were generated using the cellranger multi pipeline and imported into Seurat v5.2.1^9^ for preprocessing. Nuclei with ≤200 detected genes or ≥0.25% mitochondrial reads were excluded, and the top 1% of nuclei per library with abnormally high UMI counts were removed to eliminate potential multiplets. Doublets were identified using DoubletFinder v2.0.6^10^ with expected doublet rates derived from 10x recovery estimates and the SCTransform normalization flag enabled. Three libraries (samples 982, 985, 1005) initially flagged for elevated mitochondrial content were retained after manual inspection of their expression profiles. Prior to normalization, Apoe and App were removed from the count matrices to prevent alignment bias due to strain-specific sequence differences in knock-in genotypes; raw counts for these genes were retained in metadata for visualization only. Normalization and variance stabilization were performed using sctransform v2^11^. UMAP embeddings generated using 15–30 principal components showed stable cluster structure, and 15 PCs were selected for downstream analysis. No batch correction was applied, as no significant batch effects were detected.

##### Cluster resolution optimization

The optimal clustering resolution was determined using a Random Forest^12^–based optimization approach with silhouette score evaluation and subject-wise cross-validation across a range of resolution parameters (0.02–1.2), as described by George et al. (2022)^13^. and implemented in the clustOpt R package^14^.

##### Clustering and cell-type annotation

Graph-based clustering was performed in Seurat using *FindNeighbors* and *FindClusters* (Louvain algorithm) with 15 principal components and a resolution of 0.04, resulting in seven biologically distinct clusters. Marker genes were identified using *FindAllMarkers* (Wilcoxon rank-sum test, minimum expression = 25%). Cell types were assigned with the ScType framework^15^, referencing three curated mouse marker databases: CellMarker 2.0 brain-specific markers, custom cortex markers, and canonical hippocampal gene sets^16^. Final annotations were manually refined by integrating ScType predictions with *FindAllMarkers* output.

##### Association of cluster membership with genotype

Associations between cell-type composition and genotype were evaluated using the LODopt R package^17^, which estimates changes in absolute cluster composition while accounting for the compositional structure of single-cell data through log-odds ratio optimization. Cluster counts were aggregated by sample, and genotype was modeled as a fixed effect (wild-type as reference) within a generalized linear mixed-effects model framework. Normalization factors were optimized to minimize bias in log-odds ratio estimation, and statistical significance was assessed at *p* < 0.05.

#### WGCNA data preprocessing and module construction

For the snRNA-seq datasets, single-cell counts were aggregated to pseudobulk profiles by genotype, treating each biological sample ID as a replicate. Pseudobulk aggregation was performed with hdWGCNA^18^ to generate sample-level expression matrices. From this point onward, all analyses used the WGCNA R package^19^. Raw counts were normalized to log2 counts per million (log2CPM). To remove low-quality and uninformative features, we excluded housekeeping genes (ribosomal, mitochondrial, histone related), filtered out genes below the 10th percentile of median absolute deviation, and retained the top 5,000 most variable genes among the remainder.

#### Cell types and WGCNA

WGCNA was performed separately for four major cell types identified by clustering: Cluster 0 (pyramidal neurons), Cluster 1 (astrocytes), Cluster 3 (interneurons), and Cluster 5 (microglia). The soft-thresholding power was chosen to approximate scale-free topology and set to 24 (pyramidal), 14 (astrocytes), 14 (interneurons), and 7 (microglia). Networks were built with ConstructNetwork using cell-type–specific parameters: mergeCutHeight = 0.30, deepSplit = 1 (pyramidal); mergeCutHeight = 0.35, deepSplit = 1 (astrocytes); mergeCutHeight = 0.15, deepSplit = 1 (interneurons); and mergeCutHeight = 0.35 (microglia). Modules were detected from the topological overlap matrix (TOM) via hierarchical clustering followed by dynamic tree cutting, and dendrograms were visually inspected to confirm separation and coherence.

#### Module eigengene and hub-gene analysis

Module eigengenes (MEs) were computed as the first principal component of each module’s expression matrix, and gene–module membership (kME) was defined as the Pearson correlation between each gene and its corresponding ME. To standardize polarity across modules, ME signs were aligned such that the correlation between each ME and the mean expression of the top 50 hub genes (ranked by |kME|) was positive; when negative, both ME and kME values were inverted. MEs were then WT-centered by subtracting the WT group mean and dividing by the WT standard deviation, such that positive and negative values represent relative increases or decreases in module activity compared to WT. Hub genes were defined as the top 25 genes per module ranked by |kME|, and these hub sets were used to compute single-cell module activity scores using Seurat’s AddModuleScore. Module activity was subsequently summarized at the sample and cell-type levels for downstream genotype comparisons.

#### Contribution analysis

For contribution analysis, WT-centered MEs were used to quantify genotype-specific shifts in module activity. For each module, the magnitude of activity change in each genotype group was calculated as the absolute deviation from the WT module eigengene mean. These absolute shifts were then summed across all modules within a given cell class to obtain the total module-level perturbation attributable to E4 alone, NLF alone, or their combined interaction in the E4NLF genotype. The resulting three totals represent the overall transcriptional disruption associated with each factor, and were converted to proportional contributions by dividing each total by the sum across all three groups. These relative contributions were visualized as pie charts to illustrate the dominant driver(s) of co-expression network dysregulation within each cell type. Contribution scores (cE4, cNLF, cE4×NLF) were compared between neuronal (interneuron and pyramidal) and glial (microglia and astrocyte) modules using the Wilcoxon rank-sum test (Mann–Whitney U, non-parametric, two-sided). Multiple comparisons were corrected using the Bonferroni method.

#### Cytoscape network export

For each module, pairwise Pearson correlations among member genes were calculated, and edges with |r| ≥ 0.3 were retained to define gene–gene connectivity networks. Node attribute tables included gene identifiers and their corresponding kME values. Edge and node tables were exported and visualized in Cytoscape v3.9.1^20^. For visualizations, node size was scaled according to hubness, defined as the absolute value of kME, such that larger nodes represent genes with stronger intramodular connectivity. Node color reflected the signed kME value, allowing positive and negative associations with the module eigengene to be distinguished.

#### Genotype × module ME heatmap

For each module, sample-level module eigengenes (MEs) were extracted from the pseudobulk WGCNA output. Genotype identity was parsed from sample IDs and appended to the ME table. To compare module activity across genotypes, we computed WT-centered z scores for each module by subtracting the WT mean ME and dividing by the pooled standard deviation across genotypes, providing an effect-size–scaled representation of ME shifts. To statistically assess genotype-dependent module changes, we performed limma contrasts (genotype vs. WT) using the module × sample ME matrix and applied Benjamini–Hochberg FDR correction (FDR < 0.05 significance threshold).

#### Module wise gene heatmaps

For each module, genes present in the pseudobulk matrix were ranked by absolute kME, with variance as a fallback when kME was unavailable. Up to the top 100 genes were retained. Gene level WT centered z scores were computed using the same pooled SD approach applied gene wise, then averaged within each genotype to produce gene × genotype matrices. We plotted two views per module, the WT centered values and a row scaled version. All matrices and plots were saved alongside per contrast limma outputs.

#### Gene-level two-way ANOVA (E4, NLF, and interaction) with module-level aggregation

Two-way genotype effects were assessed at the gene level using pseudobulk log₂CPM values. Genotype was decomposed into E4 (present in E4 and E4NLF) and NLF (present in NLF and E4NLF). Within each WGCNA module (grey excluded), we fit: y∼E4×NLF for every gene. For each term (E4, NLF, E4×NLF), gene-level P values were aggregated to the module level using Fisher’s method, and FDR was controlled across modules. Module summaries included number of genes tested, fraction nominally significant, and mean η². Per-gene statistics and module-level results were saved to CSV for downstream interpretation and figure annotation.

#### KEGG pathway enrichment and visualization

For each WGCNA module (grey excluded), module gene sets were taken from the module table and standardized to module and gene. When available, genes were ordered by absolute kME; optional filters on kME threshold and top N per module were applied before enrichment. The pseudobulk expression matrix was auto oriented to genes by samples. KEGG enrichment was run per module with enrichKEGG^21^ using the species code (mmu), Benjamini–Hochberg for p value adjustment, q value cutoff 0.05, and p value cutoff 0.1. A minimum of four mapped genes per module was required to test. When available, results were converted to readable symbols with setReadable. Full and per module KEGG tables were written to CSV. Dot plots were generated per module using a fixed GeneRatio, computed as Count divided by the pathway size parsed from BgRatio. Up to 15 pathways per module were shown, ordered by adjusted p value and Count, and saved as PDF. For significant pathways (adjusted p < 0.05), pathway member genes were collected and intersected with the expression matrix. Raw log2CPM values and WT centered values were exported to Excel. WT centering was performed gene wise by subtracting the WT mean across WT samples and dividing by the gene wise standard deviation across all samples, with zero variance guarded to one. For each pathway sheet, the table included raw values per sample, WT centered values per sample, and genotype means for both raw and WT centered matrices.

## Code availability

All custom R and shell scripts used for post-Cell Ranger analysis and WGCNA in this study and data are accessible via GitHub at https://github.com/ADNetworksPPG/JP_FJ03.

